# Auditing Biomedical Literature with Blinded AI Peer Review: Where Peer Review Works, Where It Fails, and How to Strengthen It

**DOI:** 10.64898/2026.11.02.763753

**Authors:** Sergei L. Kosakovsky Pond, Stephanie J. K. Pond, Steven Weaver, Jordan D. Zehr, Darren P. Martin, Anton Nekrutenko

## Abstract

Biomedical publishing has grown more than elevenfold since 1991, outstripping human peer review. Unblinded language models cannot audit this literature objectively because they recognize famous authors and journals from training memory. We built BARS (Blinded AI Review System), an automated referee that strips titles, author names, institutions, journal headers, and dates before grading full-text manuscripts on a 10-point scale covering methods, evidence, biology, and reproducibility. We used BARS to generate 1,874 blind reviews across 1,714 unique papers. BARS passed 100% of verified sound controls, caught 77% to 91% of internally flawed papers, eliminated a +0.35-point prestige boost, and more than doubled human referee consistency. Across 320 recent open-access articles (2020–2025) from eight fields, 54% failed on core scientific rigor (*<* 5.0*/*10; 41% excluding megajournals). Revision at selective journals improved preprints more than twice as much as revision at commercial megajournals (+0.62 vs. +0.30 points; baseline-adjusted *p* = 0.033), and BARS recalled 98% of unresolved human referee critiques across 100 open-review papers. Across 915 date-blinded papers from 1991 to 2025, average scores fell from 7.6*/*10 to 5.8*/*10 as megajournals and template database studies expanded. Blinded self-audits of 238 papers by two senior authors—and of this manuscript itself—caught real statistical, control, and table errors in our own work. At four cents per paper, BARS offers a practical quality check for authors, journals, and readers (https://veg.github.io/BARS/).

## 1 Introduction

Scientific publishing rests on peer review: before a study appears in print, independent experts check that its experiments, statistics, and conclusions hold up. Over the past 35 years, however, the volume of biomedical papers indexed each year in PubMed across our core biological fields grew more than elevenfold, rising from roughly 58,000 articles in 1991 to nearly 650,000 in 2025, and nearly 2.0 million per year across all of PubMed. Open-access full-text papers grew nearly 300-fold over the same period. The pool of human peer reviewers cannot keep pace with the growth of submitted publications. Longstanding theoretical work warned that small sample size, flexible data mining, and pressure to publish positive results would make many published findings false [1]. Careful follow-up investigations have repeatedly confirmed that warning. In microbiome research, laboratory reagent contamination has been mistaken for tumor or placental bacteria [2, 3, 4]. In human genetics, subtle geographic ancestry differences were misread as natural selection on human height [5, 6, 7]. In single-cell genomics, treating thousands of cells from just two or three patients as independent samples has produced waves of false discoveries [8, 9, 10].

Catching these errors after publication takes years of detective work by specialized teams. Discovering that human DNA contamination and database errors created false microbial signatures across dozens of cancer types required manual re-analysis of terabytes of raw sequencing reads [11, 12]. Showing that a faulty genotyping probe caused a spurious link between the human *CCR5* -Δ32 mutation and mortality in the UK Biobank required digging into raw chip intensities [13, 14]. Everyday peer review rarely catches such problems because human referees often disagree—showing consistency scores of just 0.20 to 0.45 on a 0-to-1 scale [15, 16]—and are heavily swayed by reputation. Controlled experiments show just how strong that prestige bias is [35, 36]. When over 3,300 referees reviewed the exact same manuscript, only 23% recommended rejection when a Nobel laureate’s name was on the cover page, compared to 48% when the paper was anonymized and 65% when attributed to an unknown junior author [36]. When a manuscript carries famous names or appears in a flagship journal, human readers and unblinded AI tools alike give it the benefit of the doubt, overlooking missing controls or math errors in supplementary tables.

Previous efforts to automate scientific quality control have taken two paths, each with a major blind spot. The first path uses rule-based software to check specific reporting formulas. In psychology, the `statcheck` program scanned over 30,000 articles and found that half had at least one *p*-value that did not match its reported test statistic [37], while the `GRIM` test caught impossible sample averages in half of small-sample studies [38]. In biomedicine, automated preprint screens such as `ScreenIT` flag missing reagent identifiers, bar graphs, or data statements [39]. While valuable, rule-based scripts only check surface syntax; they cannot tell whether a study forgot a negative extraction control or mistook batch effects for biology. The second path uses machine learning and large language models (LLMs) to mimic human peer review [40, 41, 43]. A landmark study tested GPT-4 on over 3,000 *Nature*-family papers and 1,700 machine-learning submissions, finding that GPT-4 raised many of the same points as human referees (*∼* 31%–39% overlap, matching human-to-human agreement) and that 82% of surveyed authors found the AI feedback more helpful than at least some human reviews [41]. More recently, a benchmark of 15 LLMs on randomized clinical trials seeded with 14 artificial methodological flaws (modeled on classic British Medical Journal referee trials [42]) showed that LLMs caught 46% to 93% of planted flaws compared to just 27% for historical human reviewers [43]. Yet simply pasting a published paper into an uncalibrated chatbot fails as an objective audit for two reasons. First, if the AI can see the title, authors, journal, or workspace hints, it can recall from its training memory whether a paper was praised, cited, or retracted—or recall famous error checklists—and inherit human prestige biases [40]. Second, without a calibrated scoring scale anchored against verified sound and flawed papers, open-ended “find flaws” prompts create a severe false-alarm trap: in the clinical-trial benchmark [43], the models that caught the most planted errors also falsely flagged 25% to 29% of those same error categories on clean, unmodified control trials (*r* = *−*0.77).

Here we introduce BARS (Blinded AI Review System), a strictly blinded peer-review pipeline powered by `gemini-4-argon` with internet search turned off. Before BARS reads a manuscript, an automated scrubber strips away the title, author names, universities, journal name, publication year, web links, and any retraction notices. BARS must judge the paper solely on its text, equations, figures, and tables across four plain-English questions totaling 10 points: Are the methods and statistics sound (3.5 points)? Do the data actually support the claims (2.5 points)? Are the assumptions biologically realistic (2.0 points)? Are the data and code openly shared (2.0 points)? For historical papers published before open-data mandates, we also track the 8-point Scientific Core covering just the first three questions. Across 1,874 full-text reviews of 1,714 unique papers spanning six experiments (Table 1), our investigation yields six take-home messages for scientists, editors, funders, and readers.

**Table 1:**
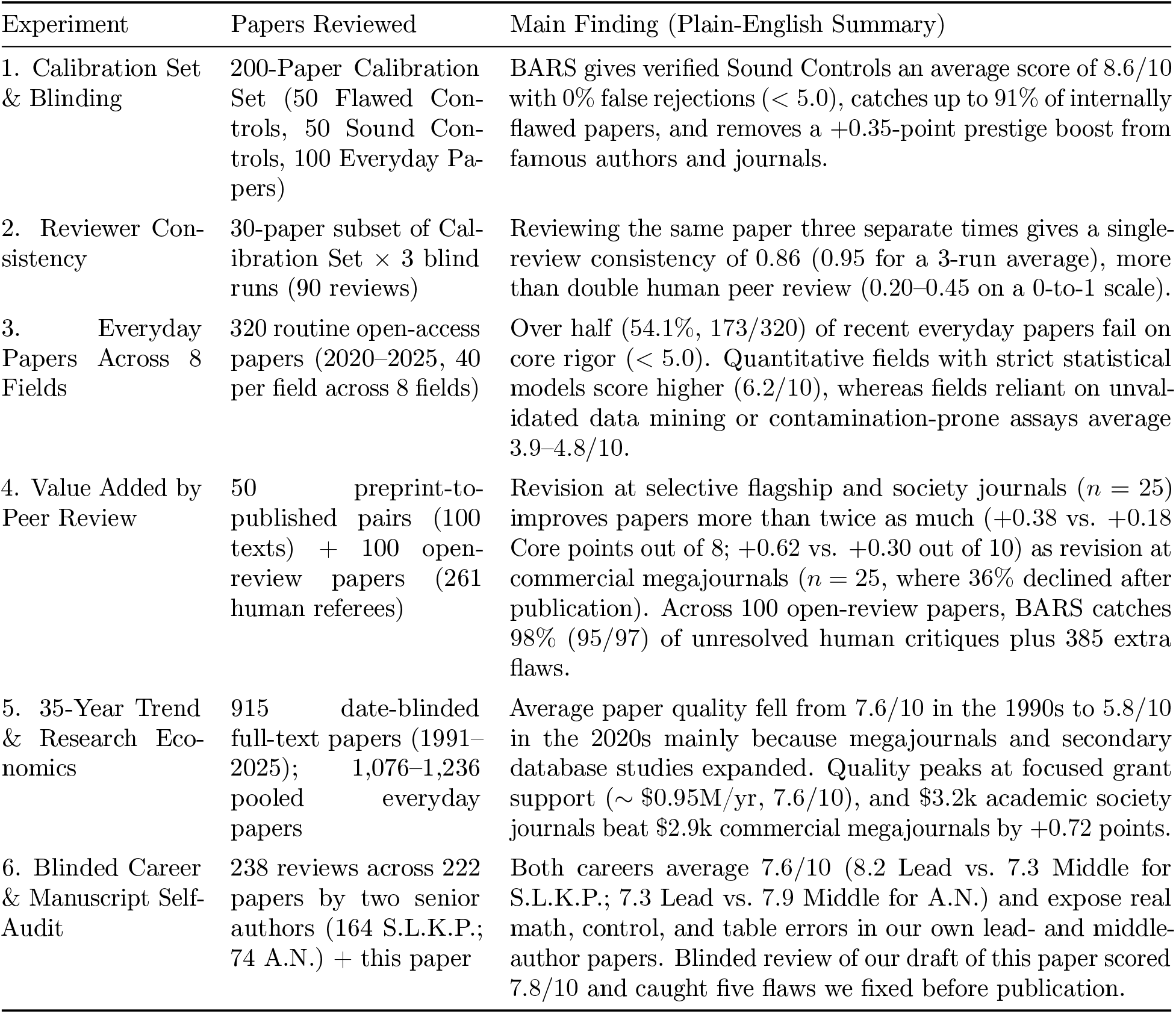
Summary of the six experiments using BARS (Blinded AI Review System; 1, 874 full-text reviews across 1, 714 unique papers). Every paper is graded without author, journal, or date metadata on a 10-point quality scale (where a score below 5.0 indicates fundamental methodological or statistical flaws) and an 8-point Scientific Core scale (which excludes open-data links). Full tables appear in the Supplementary Information (Tables S1–S7), and all reviews are searchable at https://veg.github.io/BARS/.

| Experiment | Papers Reviewed | Main Finding (Plain-English Summary) |
|---|---|---|
| 1. Calibration Set & Blinding | 200-Paper Calibration Set (50 Flawed Con-trols, 50 Sound Con-trols, 100 Everyday Pa-pers) | BARS gives verified Sound Controls an average score of 8.6/10 with 0% false rejections (< 5.0), catches up to 91% of internally flawed papers, and removes a +0.35-point prestige boost from famous authors and journals. |
| 2. Reviewer Con-sistency | 30-paper subset of Cal-ibration Set × 3 blind runs (90 reviews) | Reviewing the same paper three separate times gives a single-review consistency of 0.86 (0.95 for a 3-run average), more than double human peer review (0.20–0.45 on a 0-to-1 scale). |
| 3. Everyday Papers Across 8 Fields | 320 routine open-access papers (2020–2025, 40 per field across 8 fields) | Over half (54.1%, 173/320) of recent everyday papers fail on core rigor (< 5.0). Quantitative fields with strict statistical models score higher (6.2/10), whereas fields reliant on unval-idated data mining or contamination-prone assays average 3.9–4.8/10. |
| 4. Value Added by Peer Review | 50 preprint-to-published pairs (100 texts) + 100 open-review papers (261 human referees) | Revision at selective flagship and society journals (*n* = 25) improves papers more than twice as much (+0.38 vs. +0.18 Core points out of 8; +0.62 vs. +0.30 out of 10) as revision at commercial megajournals (*n* = 25, where 36% declined after publication). Across 100 open-review papers, BARS catches 98% (95/97) of unresolved human critiques plus 385 extra flaws. |
| 5. 35-Year Trend & Research Eco-nomics | 915 date-blinded full-text papers (1991–2025); 1,076–1,236 pooled everyday papers | Average paper quality fell from 7.6/10 in the 1990s to 5.8/10 in the 2020s mainly because megajournals and secondary database studies expanded. Quality peaks at focused grant support (∼ \$0.95M/yr, 7.6/10), and \$3.2k academic society journals beat \$2.9k commercial megajournals by +0.72 points. |
| 6. Blinded Career & Manuscript Self-Audit | 238 reviews across 222 papers by two senior authors (164 S.L.K.P.; 74 A.N.) + this paper | Both careers average 7.6/10 (8.2 Lead vs. 7.3 Middle for S.L.K.P.; 7.3 Lead vs. 7.9 Middle for A.N.) and expose real math, control, and table errors in our own lead- and middle-author papers. Blinded review of our draft of this paper scored 7.8/10 and caught five flaws we fixed before publication. |

## 2 Results

### 2.1 Take-Home 1: BARS Separates Sound Science From Flawed Papers and Removes Prestige Bias

Before using BARS to judge everyday papers, we must test two things: whether it falsely penalizes solid science, and whether it reliably catches flawed methods, invalid statistics, and unsupported conclusions without seeing who wrote the paper. Importantly, we specified and froze our four-category, 10-point grading rubric (`REVIEW_TEMPLATE.md`, commit `01665db`) *before* sampling or evaluating any manuscripts, ensuring that the metrics were not tuned post hoc to fit the papers. To test the frozen rubric, we assembled a 200-Paper Calibration Set across five core biomedical fields: viral evolution, host-associated and cancer microbiome studies, human statistical genetics, single-cell gene expression, and molecular evolution. The 200-Paper Calibration Set consists of three groups:

1. **Flawed Controls (Cohort A, 50 papers, 10 per field):** Cohort A consists of published papers that were formally retracted or disproven by later studies because of severe methodological or statistical errors [17, 18, 19, 11, 20, 21, 13, 22, 23, 24].
2. **Sound Controls (Cohort B, 50 papers, 10 per field):** Cohort B consists of verified high-rigor studies—independently validated empirical biological discoveries and at most two foundational methods per field [25, 12, 5, 6, 28, 26, 27, 29].
3. **Everyday Papers (Cohort C, 100 papers, 20 per field):** Cohort C consists of routine open-access research articles published between 2020 and 2025, sampled from Europe PubMed Central without filtering for journal prestige or citation count (also referred to as “wild” literature in our supplementary tables).

BARS strips all identifying information—article titles, author names, institutions, journal names, and retraction notices—from every paper as its first step. We measured how much this blinding step changes scores by running BARS both with and without identifying headers on a 75-paper pilot subset of the 200-Paper Calibration Set (25 Flawed Controls, 25 Sound Controls, and 25 Everyday Papers) before locking our scoring cutoffs for the remaining 125 held-out papers of the Calibration Set. On the 25 Everyday Papers, hiding author and journal names lowered average scores by 0.35 points (from 5.92 down to 5.57 out of 10), showing that famous institutions and journal headers buy a 0.35-point prestige halo when unblinded. Conversely, on the 25 Flawed Controls, hiding retraction banners raised average scores from 3.46 to 4.65: when an unblinded model sees the title of a retracted paper, it recalls the retraction from training memory, whereas under blind review BARS must find the methodological flaws directly from the text and tables.

Under blind review across the full 200-Paper Calibration Set, BARS separated Sound Controls from Flawed Controls (Figure 1A, Supplementary Table S1). Sound Controls (Cohort B) scored an average of 8.6 *±* 0.9 out of 10 (median 9.0). Not a single verified Sound Control scored below our 5.0-point failure cutoff (100% specificity, 50 out of 50 passed, and 98.0%, 50 out of 51, before reassigning one pilot paper, `PMC8135569`, whose published methodological refutation placed it in Cohort A; Supplementary Table S1), and 98% (49 out of 50) scored 6.0 or higher. Meanwhile, Flawed Controls (Cohort A) averaged just 4.4 *±* 2.0 (median 4.2, *p* = 9.2 *×* 10*^−^*^16^), yielding an overall diagnostic accuracy (area under the receiver operating characteristic curve) of 0.97 across all 100 controls, and 0.98 on the 125-paper held-out portion of the Calibration Set. In 95% of the Flawed Controls it flagged (38 of 40), BARS pinpointed the statistical error, missing control, or contamination artifact described in the published retraction notice or refutation paper.

**Figure 1:**
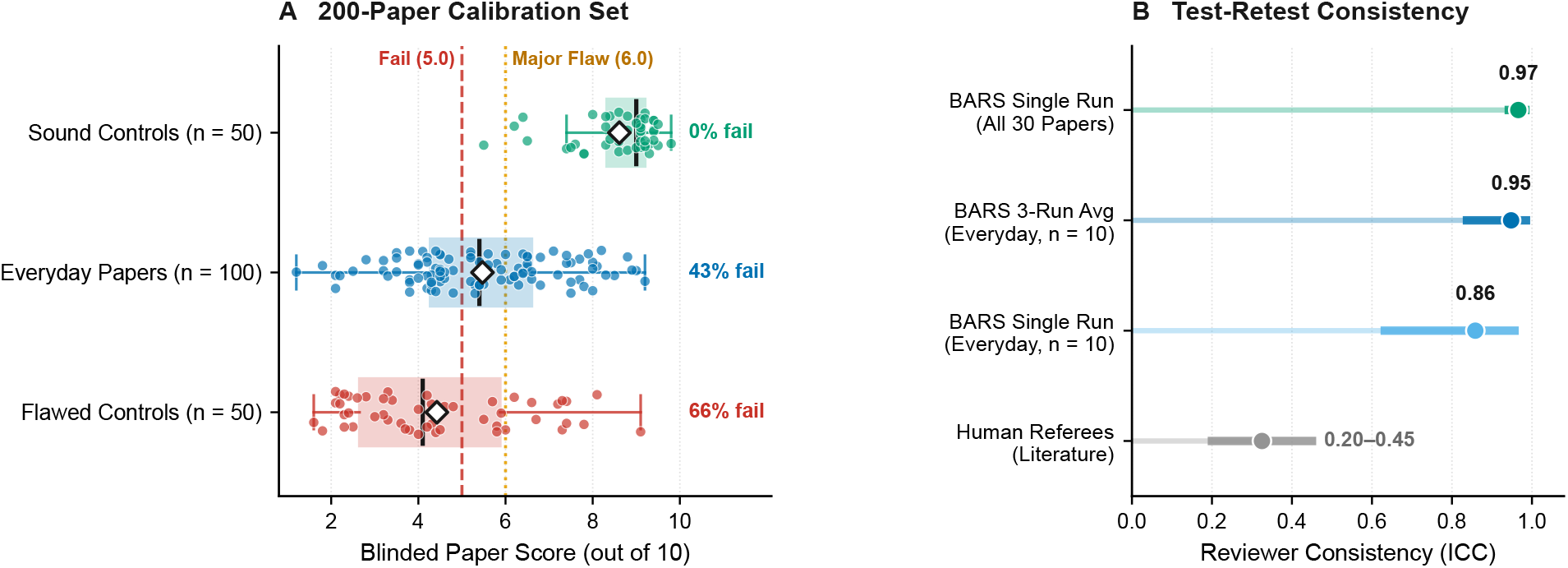
BARS separates Sound Controls from Flawed Controls in the 200-Paper Calibration Set with high test-retest consistency. Panel A shows blinded review scores (/10) across 50 Sound Controls (Cohort B, green), 100 Everyday Papers from 2020–2025 (Cohort C, blue), and 50 Flawed Controls that were retracted or refuted (Cohort A, red). Vertical lines mark the *<* 5.0 failure cutoff (red dashed line, 0% false rejections among Sound Controls) and the *<* 6.0 major-flaw cutoff (orange dotted line). Panel B shows test-retest consistency across a 30-paper subset of the Calibration Set (10 Sound Controls, 10 Flawed Controls, and 10 Everyday Papers from Panel A) reviewed blind three separate times in fresh sessions. Single-run consistency reached 0.86 in Everyday Papers (0.95 for a 3-review average, and 0.97 across all 30 papers), more than double typical human referee consistency (0.20–0.45).

Six of the 50 Flawed Controls had defects that originated inside external databases or simulation regimes instead of the manuscript’s own equations or tables (raw sensitivity across all 50 papers was 66.0% at *<* 5.0 and 76.0% at *<* 6.0), and five of those six scored above 7.0 under blind review. Three of those papers developed mathematical tests to detect natural selection on human height and applied them to public summary tables from the GIANT genetics consortium [23, 24, 30]. Years later, analyses of UK Biobank sibling data showed that the upstream GIANT summary tables contained hidden geographic bias [5, 6]. The other three papers relied on external clinical archives, microbial databases, or macroevolutionary likelihood models whose flaws only appeared under outside re-analysis or simulation [31, 32]. A reviewer reading a manuscript text alone cannot easily detect hidden biases inside an external consortium’s database. When we set aside those 5 to 6 upstream-database cases (*N_A_* = 44–45), BARS catches 76% to 77% of internally flawed papers at the strict *<* 5.0 cutoff, 89% to 91% at *<* 6.0, and 96% to 98% at *<* 6.5, with a diagnostic accuracy of 0.99 (Supplementary Table S1).

We also tested score stability across repeated runs on a 30-paper subset of the Calibration Set (10 Sound Controls, 10 Flawed Controls, and 10 Everyday Papers), reviewing each paper blind three separate times under fresh anonymous IDs (90 reviews total, Figure 1B). Across the 10 Everyday Papers, single-review test-retest consistency was 0.86 on a 0-to-1 scale (with no drift across waves, *F* = 0.37*, p* = 0.69), rising to 0.95 when averaging three runs (and 0.97 across all 30 papers from the three groups in Figure 1A). Whereas human peer-review studies measure agreement across different individuals (0.20 to 0.45 [15, 16]), BARS applies a single standardized yardstick with high run-to-run stability. An unplanned natural check supported this stability: 29 papers that were independently sampled into two different experiments under separate anonymous IDs correlated at *r* = 0.95 (*p* = 2.6 *×* 10*^−^*^15^, median score difference 0.60 points; Supplementary Table S2).

As an independent cross-check, we also replicated the recent 15-model clinical-trial benchmark of Brian et al. [43]. Because their study tested Google’s lightweight `Gemini 3.8 Flash` model and omitted a flagship Pro model, we ran BARS (`gemini-4-argon` / `Gemini 3.1 Pro`) on three matched pairs of their 2026 *Nature Medicine* randomized trials (6 blinded manuscripts total; Supplementary Section S4, Supplementary Table S7). Each pair compared an unmodified control trial against its 14-error seeded counterpart. On the error-seeded trials, BARS caught 84% of planted major flaws, including 100% of planted randomization, intention-to-treat, unvalidated-scale, and numerical-contradiction errors. At the same time, it kept false alarms on the unmodified control trials down to 2.4% (97.6% specificity vs. 71%–86% for the highest-sensitivity models in Brian et al. [43]). That yielded a net diagnostic accuracy of Youden’s *J* = 0.82, comparable to the top model in their study (`GPT-6.1 Sol` at *J* = 0.79) within the precision of an illustrative *n* = 3 trial-pair cross-check. Our calibrated 10-point score separated the unmodified trials (6.9 *±* 0.6, 100% passed) from the error-seeded versions (3.0 *±* 0.5, 100% failed below 5.0) by nearly four points with zero overlap. Notably, BARS also caught a real, unplanted flaw in a published *Nature Medicine* trial—a registered primary-outcome variance promised in the Abstract and Methods but omitted from the Results—that human peer review had missed.

### 2.2 Take-Home 2: Roughly Half of Recent Papers Fail on Scientific Rigor, With Wide Gaps Across Fields

We next audited 320 routine final published Everyday Papers (2020–2025) across eight scientific fields (*n* = 40 per field; Figure 2, Supplementary Table S2). Two hundred of these papers came from our five core biomedical fields (*n* = 40 per field, expanding the initial 100 Everyday Papers of Cohort C), and 120 came from three additional out-of-distribution fields never used during calibration: ecology and macroevolution, neuroscience and brain imaging, and AI-driven drug discovery (*n* = 40 per field). To avoid cherry-picking, all 320 papers were selected programmatically via Europe PubMed Central API queries (`OPEN_ACCESS:y AND PUB_YEAR:[2020 TO 2025]`) using fixed field-specific search terms, taking the first 40 full-text research articles per field after excluding reviews, editorials, and any paper in Cohort A or B (Methods).

**Figure 2:**
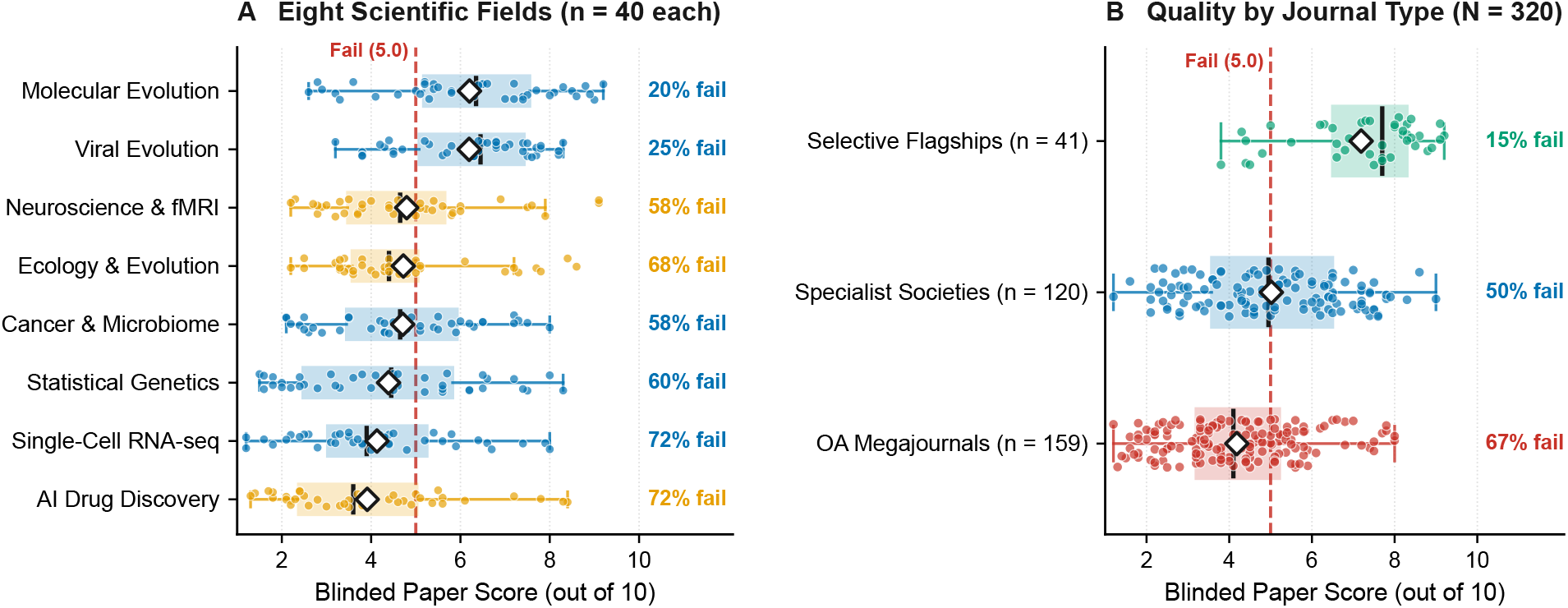
Blinded review of 320 final published Everyday Papers (2020–2025) across eight fields reveals high failure rates and a wide gap across journal types. Panel A shows BARS score distributions (/10) across five core biomedical fields (blue, *n* = 40 each) and three additional out-of-distribution fields (orange, *n* = 40 each). Quantitative fields with explicit null models (Molecular Evolution and Viral Evolution, 6.2*/*10, 20%–25% failed) score substantially higher than fields with frequent unvalidated data-mining studies or uncalibrated assays (Single-Cell Transcriptomics, 4.1*/*10, and AI Drug Discovery, 3.9*/*10, both 72.5% failed). Panel B groups the same 320 published papers by journal type: selective flagship journals average 7.2*/*10 (15% failed), specialist society journals average 5.0*/*10 (50% failed), and open-access megajournals average 4.2*/*10 (67% failed).

Across all 320 Everyday Papers combined, the average score was 4.88 *±* 1.92 out of 10 (median 4.70, Scientific Core 3.97 *±* 1.67*/*8.0), and 54.1% (173*/*320) scored below the 5.0-point failure cutoff (70.6% scored below 6.0, and only 6.9% reached 8.0 or above). Across the 200 papers in our five core fields, the average score was 5.12 *±* 1.96 (47.0% failed below 5.0; 43.0% in the initial 100-paper Cohort C subset). Across the 120 papers in the three out-of-distribution fields, the average score was 4.48 *±* 1.79, with 65.8% (79*/*120) failing below 5.0 (82.5% below 6.0). Adjusting for BARS sensitivity and specificity at the *<* 5.0 cutoff via the Rogan–Gladen estimator implies that roughly 70% (95% bootstrap CI: 58%–86%; or 69%–70% even if wild specificity is 90%–95%) of recent open-access papers in these fields contain fundamental methodological flaws that undermine their main conclusions.

Paper quality varied substantially across the eight fields (*n* = 40 per field; Figure 2A). Quantitative fields that require explicit mathematical null models performed best: molecular evolution averaged 6.20 *±* 1.88 (8 of 40 failed below 5.0, or 20.0%), and viral evolution averaged 6.19 *±* 1.44 (10 of 40 failed, 25.0%). Moderate average scores occurred in neuroscience and brain imaging (4.80 *±* 1.75, 23 of 40 failed, 57.5%, mostly small-sample functional MRI or clinical correlation studies) and ecology and macroevolution (4.72 *±* 1.62, 27 of 40 failed, 67.5%). Host-associated and cancer microbiome studies—balanced across exactly 20 oncology microbiome studies (4.46 *±* 1.54, 60.0% failed) and 20 host-associated dysbiosis studies (4.96 *±* 1.79, 55.0% failed)—averaged 4.71 *±* 1.67 overall (23 of 40 failed, 57.5%, most commonly cross-sectional 16S surveys lacking negative extraction controls or compositional statistics). Statistical genetics averaged 4.39 *±* 2.00 (24 of 40 failed, 60.0%), where failing studies were dominated by two-sample Mendelian randomization templates using weak or pleiotropic instruments and genome-wide association studies with circular marker selection. Single-cell transcriptomics averaged 4.13 *±* 1.76 (29 of 40 failed, 72.5%), driven by multi-gene cancer prognostic signatures lacking independent validation (including `FEXP_ANON_096`, 1.2*/*10, which retained unedited LLM placeholders `“e.g., GeneA, GeneB, GeneC“` and a copy-pasted “bladder cancer” methods paragraph inside an esophageal carcinoma paper) and pseudoreplicated single-cell studies. The lowest average appeared in AI drug discovery and structural bioinformatics (3.91 *±* 1.89, 29 of 40 failed, 72.5%), where failing papers routinely claimed therapeutic drug hits from computer docking scores without experimental binding assays.

Scores of these 320 final published articles also tracked closely with journal selectivity and downstream citations (Figure 2B). To test how publishing models relate to rigor, we grouped journals into three categories. *Selective flagship journals* are multidisciplinary or top-tier journals with strict editorial screening and low acceptance rates (such as *Nature*, *New England Journal of Medicine*, *PNAS*, and *eLife*). *Specialist society journals* are field-specific journals run by academic societies or university presses (such as *Molecular Biology and Evolution*, *Journal of Virology*, and *Genetics*). *Open-access megajournals* are high-volume commercial open-access journals that charge an article processing fee and instruct referees to evaluate only technical soundness without judging perceived importance (such as *PLOS ONE*, *Scientific Reports*, *Frontiers*, and *MDPI* series). Across all 320 Everyday Papers, final articles published in selective flagship journals (*n* = 41) averaged 7.19 *±* 1.52 (median 7.70) with only 14.6% (6*/*41) failing below 5.0. Final articles in specialist society journals (*n* = 120) averaged 5.02 *±* 1.81 (median 4.95, 50.0% failed). Final articles in open-access megajournals (*n* = 159) averaged just 4.18 *±* 1.58 (median 4.10), with 67.3% (107*/*159) failing below 5.0. Even when all 159 megajournal articles are excluded, 41.0% (66*/*161) of articles in flagship and specialist society journals still fail below 5.0 (50.0%, 60*/*120, in society journals alone). Because BARS was blinded to journal identity, the 3.01-point gap between flagship journals and megajournals reflects real differences in experimental design and statistical care.

### 2.3 Take-Home 3: Peer Review Substantially Improves Papers at Selective Journals, but Not at Megajournals

Do selective journals score higher simply because better papers are submitted to them, or does rigorous peer review actually improve manuscripts before publication? To measure the direct impact of peer review, we collected 50 matched pairs of initial version-1 preprints (from bioRxiv, medRxiv, and Research Square) and their final peer-reviewed journal versions. Half (*n* = 25) were published in selective flagship or society journals (such as *Nature*, *PLOS Biology*, *eLife*, *Nucleic Acids Research*, and *PLOS Pathogens*), and half (*n* = 25) were published in open-access megajournals (such as *PLOS ONE*, *Scientific Reports*, *Frontiers*, *PeerJ*, and *MDPI* series). We stripped all formatting, version labels, and metadata from all 100 documents and had separate BARS instances review the preprint and published version of each study (Figure 3A, Supplementary Table S3). We paired this 50-study before-and-after test with a 100-paper audit of final published open-review articles (*n* = 50 selective vs. *n* = 50 megajournal) that publicly archive their full pre-publication referee reports and author revision responses (Figure 3B). To ensure that simple formatting differences in where journals place Data Availability statements could not skew the preprint-to-published comparison, we measured improvement on both the 8-point Scientific Core (methods, claim alignment, and biology) and the 10-point overall score.

**Figure 3:**
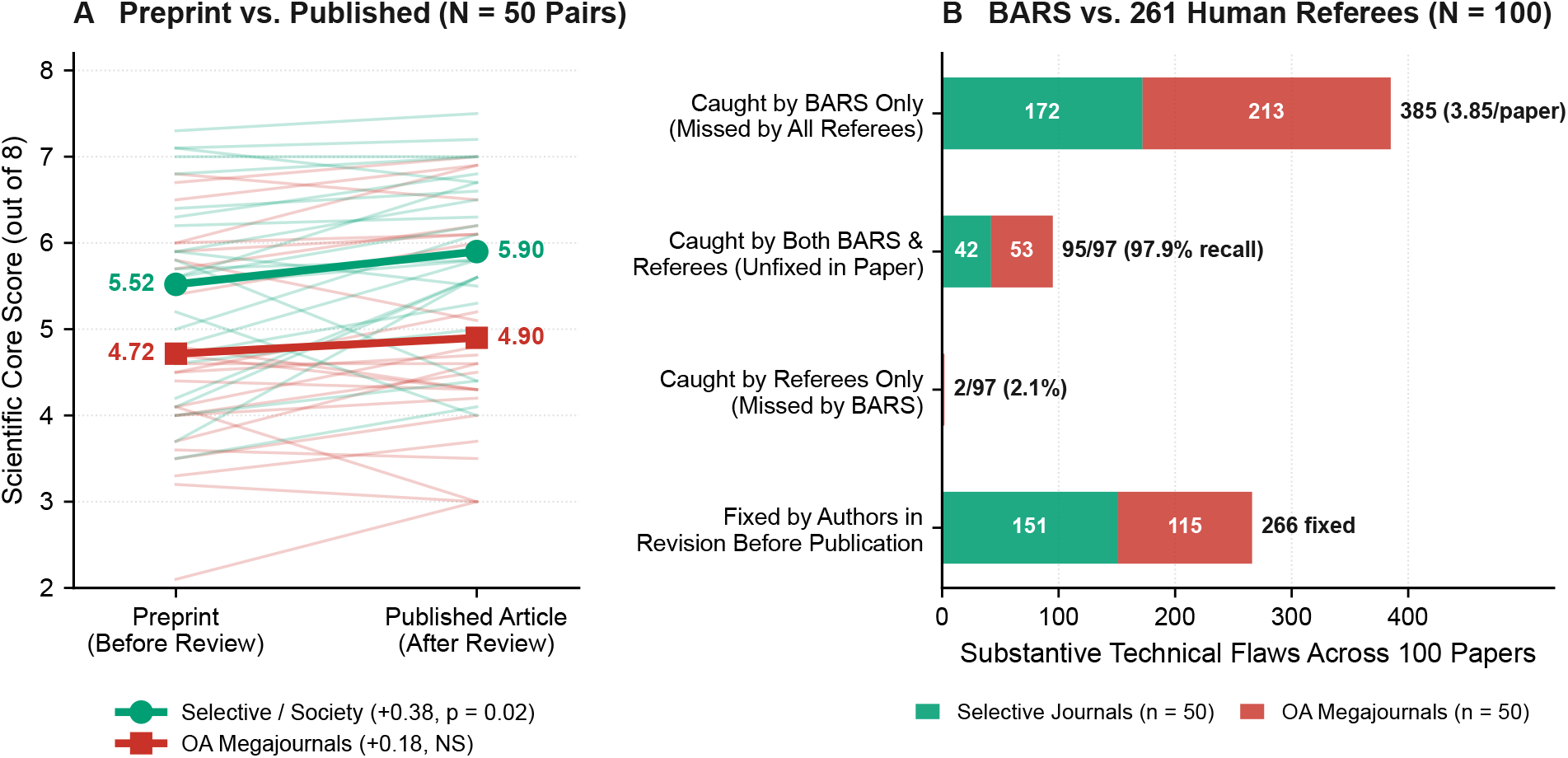
Peer review substantially improves manuscripts at selective journals, and BARS recalls 98% of unresolved human referee critiques while catching 385 additional flaws across 100 open-review papers. Panel A tracks the 8-point Scientific Core score from initial blinded Preprint (Before Review) to final blinded Published Article (After Review) across 50 matched study pairs (100 blinded texts). Studies revised at selective flagship and society journals (*n* = 25, green) gained +0.38 Core points (*p* = 0.019, and +0.62*/*10 overall, with 20 of 25 improving). By contrast, studies at open-access megajournals (*n* = 25, red) gained only +0.18 Core points (*p* = 0.089, not significant; +0.30*/*10 overall, with 9 of 25 declining after publication). Panel B compares blinded BARS reviews against 261 human referees across 100 final published open-review papers (*n* = 50 selective journals in green vs. *n* = 50 megajournals in red). The top three bars show flaws remaining in the published papers: 385 caught by BARS only (3.85/paper), 95 caught by both BARS and human referees (97.9% recall of 97 unresolved human critiques), and 2 caught by human referees only (2.1%). The bottom bar shows 266 earlier referee critiques that authors fixed during revision before publication.

Across all 50 paper pairs, peer review improved scoring on the 8-point Scientific Core by an average of +0.28 *±* 0.65 points (*p* = 0.0034, Wilcoxon *p* = 0.0019; +0.46 *±* 0.81 points on the 10-point scale, *p* = 0.00020), with 36 of the 50 papers improving after review. However, splitting the 50 pairs by journal type revealed a clear divide (Figure 3A). At selective flagship and society journals (*n* = 25), peer review raised the Scientific Core by +0.38 *±* 0.75 points (from 5.52 to 5.90 out of 8.0, paired *p* = 0.019, Wilcoxon *p* = 0.0097) and raised the 10-point score by +0.62 *±* 0.88 points (from 6.90 to 7.52, *p* = 0.0017). Twenty of the 25 selective-journal papers (80%) improved, with significant gains across Methodological Rigor (*p* = 0.0086), Claim-Evidence Alignment (*p* = 0.029), and Reproducibility (*p* = 0.00025). For example, in PAIR_22 (*Nucleic Acids Research*, rising from 5.1 to 7.2*/*10), revision scrubbed a live GitHub personal access token (github_pat_…) and unedited template placeholders (Table X, Software XX) leaked in the preprint while adding validation controls. At open-access megajournals (*n* = 25), by contrast, peer review raised the Scientific Core by only +0.18 *±* 0.52 points (from 4.72 to 4.90 out of 8.0, paired *p* = 0.089, Wilcoxon *p* = 0.076), a change that did not reach statistical significance. Megajournal review raised the 10-point score by +0.30 *±* 0.71 (*p* = 0.047), less than half the gain at selective journals. Across all 50 pairs, the unadjusted two-sample difference in raw gains was +0.20 Core points (*p* = 0.29; +0.37 points, *p* = 0.13 in our 20-pair pilot) and +0.32 total points (*p* = 0.16). Because selective preprints started nearly a full point higher (6.90 vs. 6.02) and faced ceiling compression on the 10-point scale, baseline-adjusted analysis of covariance showed that selective peer review achieved a +0.50 *±* 0.23-point higher post-review total score (*t* = 2.20*, p* = 0.033; +0.35 *±* 0.18 Core points, *p* = 0.063). In addition, 9 of 25 megajournal papers (36%) and 4 of 25 selective-journal papers (16%) scored *lower* after peer review than as preprints (Figure 3A). Inspecting the largest post-review drops pointed to two causes. Most often, authors responded to referee requests by adding new post-hoc analyses, benchmark comparisons, or summary tables that introduced fresh statistical flaws or table errors that referees never re-checked before acceptance. In PAIR_31 (*Nature Communications*, Core dropping from 5.2 to 4.0 out of 8.0), the revised article added 29 KB of text, including a new network analysis that mistook floating-point machine zero (EV = 9.88 *×* 10*^−^*^17^) for a biological keystone hub taxon. In `PAIR_38` (*Scientific Reports*, Core 5.8 to 5.1), the revision downsampled the reference control library to match the new assay’s 80%–90% PCR duplication bottleneck and added an unreplicated protocol comparison lacking sample sizes or test statistics. Similarly, in `PAIR_39` (*Scientific Reports*, Core 4.7 to 4.3), a newly added table reported a population maximum *C*_trough_ (7, 702 *µ*g/L) exceeding the maximum *C*_max_ (6, 351 *µ*g/L). In the remaining cases (`PAIR_30`, `PAIR_48`), expanded methods and results tables in the published text (+8.5 to + 15.8 KB) made pre-existing spatial scaling contradictions or two-stage regression flaws more apparent than in the shorter preprint, alongside normal run-to-run scoring variance (SD*_e_* = 0.46 points).

The paired preprint data also show how initial `bioRxiv` and `medRxiv` preprints compare against published journal articles. Across all 50 version-1 preprints—evaluated before any journal peer review—BARS scores averaged 6.46 *±* 1.40 out of 10 (5.12*/*8.0 Core, with only 14.0% failing below 5.0), and preprints that later appeared in selective or society journals averaged 6.90 *±* 1.28 (5.52*/*8.0 Core, 4.0% failing). Even before peer review, those unreviewed version-1 preprints (6.46*/*10) scored higher than final peer-reviewed articles published in open-access megajournals across every cohort in our study. Specifically, published megajournal papers averaged 4.18*/*10 in our 2020–2025 everyday cohort (*p* = 3.1 *×* 10*^−^*^14^), 5.01*/*10 in our open-review cohort (*p* = 8.4 *×* 10*^−^*^6^), 5.76*/*10 across our 35-year corpus (*p* = 0.0021), and 6.33*/*10 in the paired megajournal subset. Although preprints that eventually reach publication represent a self-selected subset of all posted preprints, research groups that voluntarily post version-1 preprints on bioRxiv and medRxiv for public community scrutiny start at a substantially higher baseline of rigor than routine direct-to-megajournal submissions, and megajournal review does not close that gap.

We next compared BARS directly against human referees across a balanced cohort of 100 published open-review papers (‘2020–2025‘) whose verbatim human referee reports and editorial decision letters are publicly archived alongside the article (Figure 3B, Supplementary Table S3). Half (*n* = 50) came from selective flagship and society open-review journals (*eLife*, *PLOS Biology*, *PLOS Genetics*, *PLOS Pathogens*, and *PLOS Computational Biology*), and half (*n* = 50) came from open-review megajournals (sampled from *PLOS ONE*, the largest biomedical megajournal archiving full referee reports in Europe PMC XML). In Stage 1, BARS reviewed all 100 stripped manuscripts without access to the human peer-review files; in Stage 2, we compared the locked BARS reviews against all 261 human referee reports (363 substantive technical critiques total). We observed three main patterns.

- **A 2.5-point quality gap between selective and megajournal open review.** BARS scores averaged 7.49 *±* 1.38 out of 10 at selective open-review journals (5.87*/*8.0 Core, 8.0% failed below 5.0), where human referee reports averaged 3, 977 words. By contrast, open-review megajournal papers (*PLOS ONE*) averaged 5.01 *±* 1.69 out of 10 (3.79*/*8.0 Core, Δ = +2.48 points, *p* = 2.1 *×* 10*^−^*^12^), with 48.0% failing below 5.0 and median referee reports of 2, 462 words.
- **97.9% BARS recall of unresolved human critiques plus 385 BARS-only forensic catches.** Across the 97 substantive human referee criticisms that authors left unresolved in the final published article, a single BARS pass independently caught 95 (97.9%: 85 exact and 10 partial matches, or 87.6% strict 1-to-1 exact recall; 78.2% across all 363 human points, with 83.3% confirmed by manual human audit of five multi-round dossiers). At the same time, BARS caught 385 mathematical, statistical, and design flaws (3.85 per paper) that escaped all 261 human referees and editors. Examples included inverted odds ratios in published regression tables (`OPEN_ANON_011`, where an adjusted odds ratio of 1.2 lay outside its 95% confidence interval of 0.9–1.14) and physically impossible ELISA concentrations exceeding 100% of wet tissue weight (`OPEN_ANON_036`, 121.7 mg/mL melanin in 100 mg/mL homogenate). Other missed flaws included human cells profiled on a rodent microRNA array (`OPEN_ANON_088`) and pre-pandemic 2017–2018 cohorts used as COVID-19 outcomes (`OPEN_ANON_083`). BARS also caught *revision-induced errors* where author edits during peer review introduced impossible negative *p*-values (*p* = *−*0.006 in `OPEN_ANON_035`) or corrupted table denominators (`OPEN_ANON_086`, `OPEN_ANON_096`) that referees never re-checked.
- **Widespread human referee anomalies in megajournals (58% vs. 2%).** At selective open-review journals, 72.0% (36*/*50) of referee dossiers provided thorough technical scrutiny, 0% were superficial boilerplate, and only 1 paper (2.0%, `OPEN_ANON_015`) had a referee anomaly—an irrelevant citation demand that the *PLOS Computational Biology* editor explicitly waived. At open-review megajournals, only 12.0% (6*/*50) of dossiers provided thorough scrutiny, 38.0% (19*/*50) were superficial 3-to-6 sentence checklists, and 58.0% (29*/*50) exhibited referee anomalies. Those 29 megajournal anomalies included 19 generic or AI-generated boilerplate reviews (such as OPEN_ANON_049, where Reviewer 1 pasted raw LLM prompt instructions into Decision Letter 1, and OPEN_ANON_019, where Reviewer 2 demanded citations to indeed.com and scribbr.com blogs), 6 coercive self-citation demands (OPEN_ANON_001, OPEN_ANON_011), and 4 single-reviewer acceptances on quantitatively flawed studies.

### 2.4 Take-Home 4: Established Journals Kept Higher Scores Over 35 Years, While Overall Quality Fell in the 2020s as the Publishing Mix Shifted

Did established journals lower their peer-review standards over the past three decades, or did the mix of published studies shift as volume expanded? We sampled 1,000 open-access articles across seven 5-year windows from 1991 to 2025 and four core biomedical tracks (250 papers initially sampled per track across viral evolution, statistical genetics, gene expression, and molecular evolution). We strictly excluded 85 archival 1990s and early-2000s records whose Europe PMC XML contained only an abstract or PDF placeholder plus bibliography. That filter yielded 915 verified full-text manuscripts (226 to 233 full-text papers per track). Because Europe PMC XML stores only the main article body, equations, and inline tables—and because external supplementary files rarely existed in the 1990s—BARS graded the self-contained main manuscript text across all seven windows while using both the 10-point overall score and the 8-point Scientific Core (which removes any penalty for modern open-data repository links). We scrubbed every date, year, author name, and journal title from all 915 manuscripts and scored them blind with BARS (Figure 4A, Supplementary Table S4).

**Figure 4:**
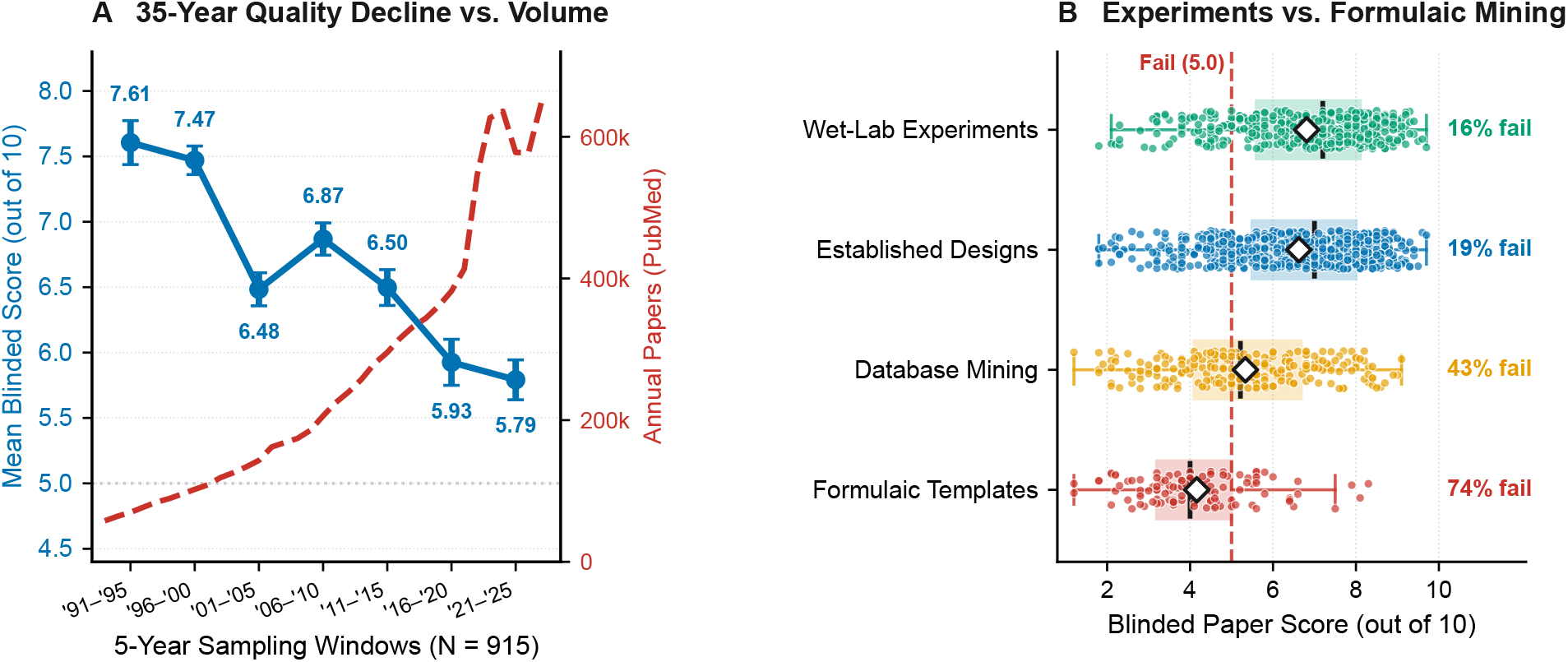
A 35-year quality decline driven by the rise of open-access megajournals, secondary database mining, and formulaic study templates. Panel A plots date-blinded BARS manuscript scores (/10, blue) across all seven 5-year sampling windows from 1991–1995 to 2021–2025 (*N* = 915 verified full-text papers), alongside the 11-fold surge in annual PubMed biomedical paper volume (red dashed curve). Scores fell in two waves: from 7.61–7.47 in the 1990s to 6.48–6.87 in the 2000s, and down to 5.93–5.79 in 2016–2025 (34.8% failed below 5.0 in the 2020s). Panel B compares study types across all 1,076 everyday full-text papers in our study: primary wet-lab experiments average 6.8*/*10 (16% failed) and established experimental or genomic designs average 6.6*/*10 (19% failed), whereas pure secondary database mining averages 5.3*/*10 (43% failed) and formulaic templates average 4.2*/*10 (74% failed).

Across the 915-paper full-text cohort, overall quality fell across two distinct waves over the 35-year period (Figure 4A). Across the seven 5-year windows, scores started high in 1991–1995 (7.61 *±* 1.43, *n* = 73, 6.8% failed below 5.0) and 1996–2000 (7.47 *±* 1.22, *n* = 124, 4.0% failed; 1991–1999 calendar-decade mean 7.63 *±* 1.25, *n* = 154, 3.9% failed). The first drop occurred in 2001–2005 (6.48 *±* 1.51, *n* = 142, 18.3% failed) before stabilizing in 2006–2010 (6.87 *±* 1.48, *n* = 144,11.8% failed; 2000–2009 calendar-decade mean 6.76 *±* 1.48, *n* = 249, 14.1% failed) and 2011–2015 (6.50 *±* 1.64, *n* = 144, 20.1% failed). A second sharp drop followed in 2016–2020 (5.93 *±* 2.11, *n* = 144, 36.1% failed) and 2021–2025 (5.79 *±* 1.83, *n* = 144, 31.9% failed; 2020–2025 calendar-decade mean 5.81 *±* 1.97, *n* = 273, 34.8% failed below 5.0 and 50.9% below 6.0). Across all 915 papers, composite quality declined by 0.58 points per decade (*r* = *−*0.33*, p* = 1.2 *×* 10*^−^*^24^; *−*0.68 scaled points/decade on the 8-point Scientific Core, and *−*0.77 points/decade when weighting all 28 calendar years equally, *p* = 6.8 *×* 10*^−^*^7^). Over 93% of that drop (*−*0.54 raw points/decade, *p* = 8.7 *×* 10*^−^*^29^) occurred inside the 8-point Scientific Core, whereas the 2-point Reproducibility score changed by only *−*0.04 points/decade (1.39*/*2.0 in the 1990s vs. 1.24*/*2.0 in the 2020s). Every subfield track declined significantly (*p ≤* 3.1 *×* 10*^−^*^6^): gene expression (*r* = *−*0.37), viral evolution (*r* = *−*0.36), statistical genetics (*r* = *−*0.31), and molecular evolution (*r* = *−*0.30). Even though BARS could usually guess a paper’s decade from the laboratory techniques described in the text (92.4% accuracy), papers that looked older did not receive higher scores than papers from the same calendar year that looked newer (partial correlation *r* = *−*0.009*, p* = 0.83), arguing against nostalgia bias in favor of older methods.

We separated within-journal changes from shifts in the publishing mix by analyzing within-journal trends across the 915-paper cohort and comparing study designs across all 1,076 everyday full-text papers in our study (Figure 4B, Supplementary Table S4).

- **Two waves of ecosystem expansion and compositional shift explain the bulk of the decline.** Across 549 papers published in 30 recurring multi-year journals, the within-journal score change was modest (*−*0.23 points/decade, *p* = 0.027), less than two-fifths of the*−*0.58-point-per-decade overall drop. A standard demographic decomposition (Oaxaca–Blinder) showed that 54.1%–58.3% (0.98 points) of the decline from the 1990s to the 2020s came from shifts in the mix of journals and study designs. In the 1990s, open-access full texts in Europe PMC came almost entirely from selective society and university press journals (such as *Journal of Experimental Medicine* and *Journal of Cell Biology*, 0% megajournals). The first score drop in 2001–2005 coincided with the launch of first-generation commercial open-access series (such as the *BMC* series in 2000, with commercial OA venues rising from 0% of the cohort in the 1990s to 33% across the 2000s). That early-2000s wave also brought a surge of underpowered candidate-gene association studies (*n* = 50–100 subjects without ancestry controls) and small-sample cDNA microarrays (*n* = 2–3 arrays without false-discovery-rate correction). The second drop in 2016–2025 was driven by high-volume commercial megajournals (*MDPI*, *Frontiers*, and *Scientific Reports*, *>* 50% of the cohort, averaging 5.80*±*1.73 vs. 6.75*±*1.79 at selective and society venues, *p* = 1.3 *×* 10*^−^*^17^) combined with secondary database mining. In a multivariable regression across all 1,076 papers, publication decade (*β* = *−*0.52/decade*, p <* 10*^−^*^15^), megajournal venue (*β* = *−*0.51*, p* = 3.0 *×* 10*^−^*^6^), and dry-lab modality (*β* = *−*0.71*, p* = 6.3 *×* 10*^−^*^10^) each remained independently significant, and restricting the 35-year cohort to non-megajournals (*n* = 493) still showed a *−*0.49-point-per-decade decline (*p* = 9.6 *×* 10*^−^*^16^).
- **Wet-lab experiments vs. pure database mining.** Primary wet-lab studies (*n* = 408) averaged 6.81 *±* 1.68 (16.4% failed) and hybrid wet-lab/computational studies (*n* = 351) averaged 6.56 *±* 1.73 (19.9% failed). By contrast, purely computational re-analyses of secondary public databases (*n* = 230) without experimental validation averaged 5.33 *±* 1.78 (42.6% failed, *p* = 5.5 *×* 10*^−^*^23^).
- **Established experimental and genomic designs vs. formulaic templates and unedited AI text.** Well-validated genomic assays (*n* = 348, 6.65 *±* 1.65, such as replicated bulk RNA-seq, deep mutational scanning, and ChIP-seq with input controls) and established experimental or statistical designs (*n* = 616, 6.62 *±* 1.71, such as biochemical binding assays, viral neutralization titers, and maximum-likelihood phylogenetics) performed consistently well. By contrast, formulaic template papers (*n* = 112)—such as fitting LASSO Cox survival signatures to public TCGA cancer cohorts without external validation, running default `Seurat` clustering on two or three patient samples, chaining automated two-sample Mendelian randomization queries, or claiming drug hits from molecular docking alone—averaged 4.16 *±* 1.42 (74.1% failed, *p* = 2.0 *×* 10*^−^*^44^). In one 2025 paper (`PMC12473078`, 1.2*/*10), BARS caught an unedited AI chatbot response and internal artifact_id UUID pasted verbatim inside the published Discussion.

### 2.5 Take-Home 5: A Blinded Self-Audit of 238 Career Papers—and of This Manuscript Itself

We next applied BARS to our own published work and to this manuscript, reporting the findings candidly. We downloaded every open-access full-text paper in PubMed Central co-authored by two of our senior authors: 164 papers by Sergei Kosakovsky Pond (S.L.K.P., 2004–2026) and 74 papers by Anton Nekrutenko (A.N., 2000–2026), totaling 238 blinded reviews across 222 unique papers (including 16 joint papers audited independently in both batches). We stripped all titles, author names, universities, journals, years, and recognizable lab software names (such as `HyPhy`, `Datamonkey`, `HIV-TRACE`, `Galaxy`, `UseGalaxy`, `Planemo`, and `DuNovo`) and had BARS grade all 238 papers blind (Figure 5, Supplementary Table S5).

**Figure 5:**
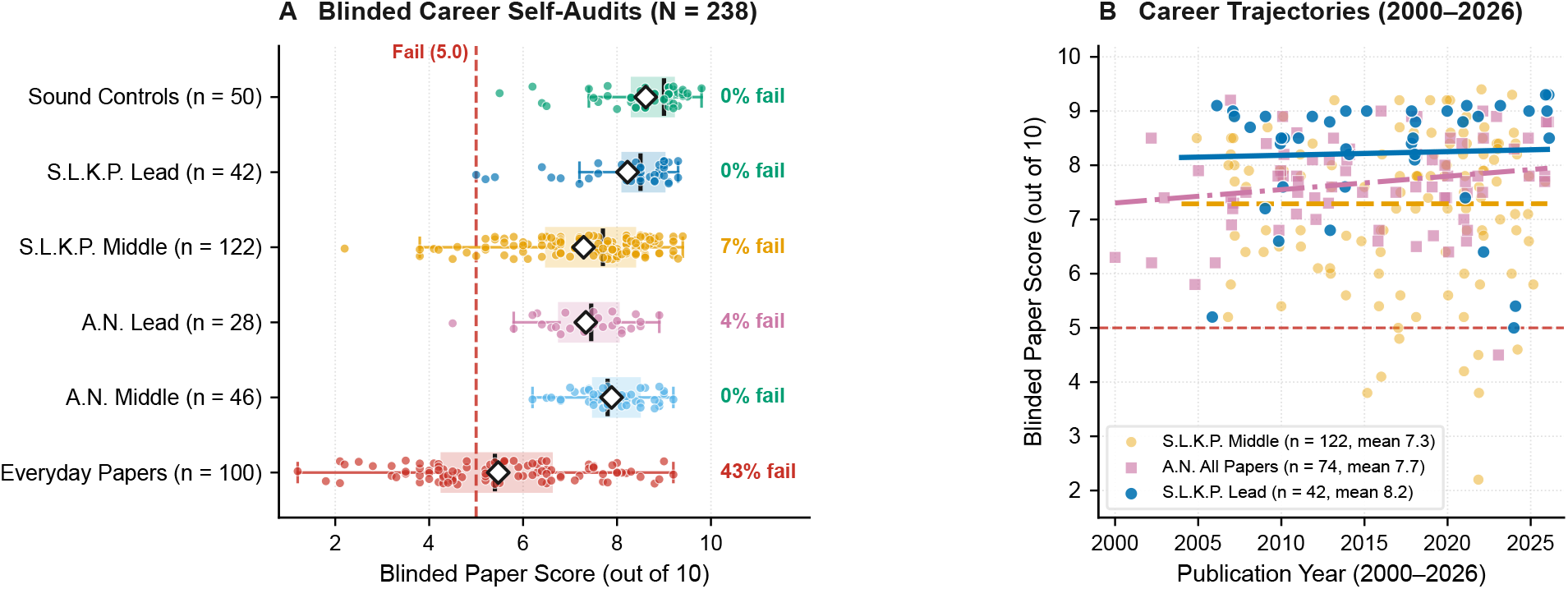
Blinded career self-audits of 238 open-access papers across two senior authors (164 by S.L.K.P., 2004–2026; 74 by A.N., 2000–2026). Panel A compares blinded BARS scores (/10) across Lead Author (First or Senior) and Middle Co-Author roles for S.L.K.P. (8.2 Lead vs. 7.3 Middle) and A.N. (7.3 Lead vs. 7.9 Middle), alongside Sound Controls (8.6*/*10) and Everyday Papers (5.5*/*10). Panel B tracks blinded scores over 26 years (2000–2026): S.L.K.P. lead-author papers stayed above 8.0, A.N. papers climbed from 7.3 in the 2000s to 7.7–7.8 in the 2010s–2020s, and S.L.K.P. middle-author service collaborations drifted downward with the broader literature.

Both careers averaged 7.6 out of 10 (7.53*±*1.38, median 7.85 for S.L.K.P.; 7.68*±*0.86, median 7.75 for A.N.), and the 16 joint papers audited independently in both batches showed strong agreement (7.82 vs. 7.88, *r* = 0.65*, p* = 0.0062, with Reproducibility scores of 1.74 vs. 1.76*/*2.0 across -only and + `<back>` XML). The two portfolios showed opposite lead-vs.-middle authorship splits, undercutting any assumption that low scores only happen when external collaborators make mistakes (Figure 5A, Supplementary Table S5). In S.L.K.P.’s portfolio, the 42 Lead Author papers averaged 8.22 *±* 1.11 (0% below 5.0, 7.1% below 6.0) versus 7.29 *±* 1.39 across 122 Middle Co-Author papers (*p* = 3.0 *×* 10*^−^*^5^). The lead-vs.-middle gap held after excluding 16 early records with unredacted github.com/veg links (+0.96 points) and within dry-lab studies (+0.84 points, *p* = 3.4 *×* 10*^−^*^4^), largely because 79% (33*/*42) of S.L.K.P.’s lead papers are foundational methods or theory studies. In A.N.’s portfolio, by contrast, the 46 Middle Co-Author papers—dominated by multi-lab `Galaxy`, Vertebrate Genomes Project, and viral surveillance consortia—averaged 7.88 *±* 0.72 (0% below 6.0). Those consortium studies outscored A.N.’s 28 Lead Author papers (7.34 *±* 0.96, *p* = 0.012, 3.6% below 5.0, 7.1% below 6.0), which included early 2000–2005 comparative genomics studies and short software notes. Over time, A.N.’s papers rose from 7.30 in the 2000s to 7.79 in the 2010s and 7.74 in the 2020s (Figure 5B).

When a paper has a flawed statistical test, uncalibrated threshold, or table bug, the error stands regardless of whether we led the study or joined as co-authors. At the same time, a low BARS score is a technical diagnosis of specific analytical or reporting flaws, not a verdict of misconduct or a call to retract the paper. If every paper with a flawed statistical test or overstated conclusion were retracted, roughly half of the published literature would vanish. Most lower-scoring studies—including our own—contribute useful software, data, or biological observations alongside technical defects that should be documented and corrected. Across our own lead-author papers, BARS caught substantive errors that had slipped past both us and journal peer review:

- **Table copy-paste errors and gap-column misclassifications in S.L.K.P. lead papers (Scores 5.0 to 5.4/10).** In a 2024 HIV network paper (`AUTO-TUNE`, 5.0*/*10, Senior Author), BARS caught identical copy-pasted network counts across two regional cohorts in Table 2, a row claiming 1,364 connected nodes from 295 sequences, and 100*×* unit mix-ups. In a 2006 HIV-1 selection paper (5.2*/*10, First Author), BARS flagged a multiple-testing base-rate fallacy (7 hits across 296 unadjusted codon tests at *p ≤* 0.05, matching the *∼* 7 false positives expected by chance). In a 2024 mammalian *Rem2* paper (5.4*/*10, Senior Author), BARS caught 251 gap-dominated alignment columns misclassified as “invariant sites” and mixed hg19/hg38 coordinates.
- **Template leakage, antibody mimicry, and table bugs in A.N. lead papers (Scores 4.5 to 6.7/10).** In a 2023 *BMC Bioinformatics* protein-interaction workflow (4.5*/*10, Senior Author), BARS caught PDB template leakage in the cross-validation benchmark and showed that the headline SARS-CoV-2 Spike–MOG receptor prediction was an antibody-mimicry artifact caused by threading MOG’s IgV domain onto anti-Spike neutralizing antibodies (`6XC2`, `7C01`). In a 2005 *PLOS Genetics* overlapping-frame study (5.8*/*10, First Author), BARS noted that claiming “evolutionary oscillation” across branches conflicted with *p* = 0.14–0.77 on every branch. In a 2002 *Genome Research K_A_/K_S_*coding-potential study (6.2*/*10, First Author), BARS noted that the 2.6% false-positive rate was a mathematical consequence of testing at *α* = 0.05 rather than an empirical specificity advantage. In a 2020 duplex-sequencing paper (`Du Novo 2.0`, 6.4*/*10, Senior Author), BARS caught Table 1 reporting 816,335 corrected families from 400,000 simulated reads and mislabeling Recall under the `Negative (%)` column. Likewise, in a 2019 runtime-prediction paper (6.7*/*10, Senior Author), BARS flagged post-execution output sizes in Table 1’s feature schema and full-dataset scaling prior to cross-validation.
- **Unchecked upstream wet-lab and framing flaws in middle-author collaborations (Scores 2.2 to 6.3/10).** Across our middle-author collaborations, lower scores arose when we analyzed downstream sequences or trajectories without auditing upstream assay controls or regression specifications. Examples include mistaking singly amplified nested-PCR carryover for 2019 Italian SARS-CoV-2 cases (2.2*/*10) [17] and misreading early viral decay during the first 6 months of therapy as ongoing HIV-1 replication alongside a negative rate table (4.1*/*10). Other examples include circular outgroup rooting and epidemic-momentum confounding in pandemic genomics (4.2 and 4.5*/*10), as well as regressing 0–24 month infant weight trajectories as the response on 24-month oral microbiota predictors (6.3*/*10).

Finally, we ran BARS on a de-identified draft of *this manuscript and its Supplementary Information* (`MANUSCRIPT_META_01`; archived verbatim at https://veg.github.io/BARS/). BARS scored our draft 7.8*/*10.0 (6.2*/*8.0 Core; Verdict: *Major Revision*) and caught five real statistical and consistency flaws that all six human co-authors had missed, each of which we fixed before publication. First, we had omitted the between-group interaction and baseline-adjusted covariance tests (+0.50 adjusted total points, *p* = 0.033; unadjusted pilot Δ = +0.37 Core points, *p* = 0.13) in the preprint-to-published experiment. Second, Europe PMC’s 5-year date queries clustered to-ward window boundaries, which prompted our equal-year-weighted regression (*−*0.77 points/decade, *p* = 6.8 *×* 10*^−^*^7^). Third, our Abstract had paired *<* 5.0 specificity (100%) with *<* 6.0 sensitivity (89%) and left 5-vs.-6 upstream-database counts unharmonized. Fourth, BARS flagged dry-lab vs. wet-lab modality confounding in the career self-audit, prompting our within-dry-lab (+0.84 points, *p* = 3.4 *×* 10*^−^*^4^) and multivariable controls, alongside fixing four minor text-vs.-table count discrepancies across cohorts. Running BARS on our own career papers and on this manuscript showed that pre-submission auditing catches table, unit, and consistency errors before editors ever see them—and that every author on a paper shares responsibility for auditing the full manuscript.

### 2.6 Take-Home 6: The Economics of Doing, Reviewing, and Publishing Science

We next examined how research budgets, journal Article Processing Charges (APCs), and review costs relate to manuscript quality by linking all papers in our study to US National Institutes of Health (NIH) grant records via Europe PMC and the NIH RePORTER API. This spanned our 200-Paper Calibration Set (Experiment 1), 320-paper Everyday Papers Across 8 Fields cohort (Experiment 3), and 915-paper 35-Year Trend cohort (Experiment 5), yielding 452 NIH-funded papers in total (222 across our 1,076-paper pooled everyday corpus). Because individual NIH grants support broad multi-paper research programs instead of single articles—and a single paper can acknowledge multiple grants—we summed the active annual award amounts of all NIH grants acknowledged by each paper during its publication year to measure the annual funding scale of the research team backing the study. To ensure that multi-paper grant sharing did not distort our conclusions, we also tested raw counts of acknowledged grants per paper (0, 1, 2–3, 4–6, and *≥* 7 grants). Finally, we mapped every journal’s open-access APC and measured token usage across all 1,874 full-text reviews of 1,714 unique papers (Figure 6, Supplementary Table S6).

**Figure 6:**
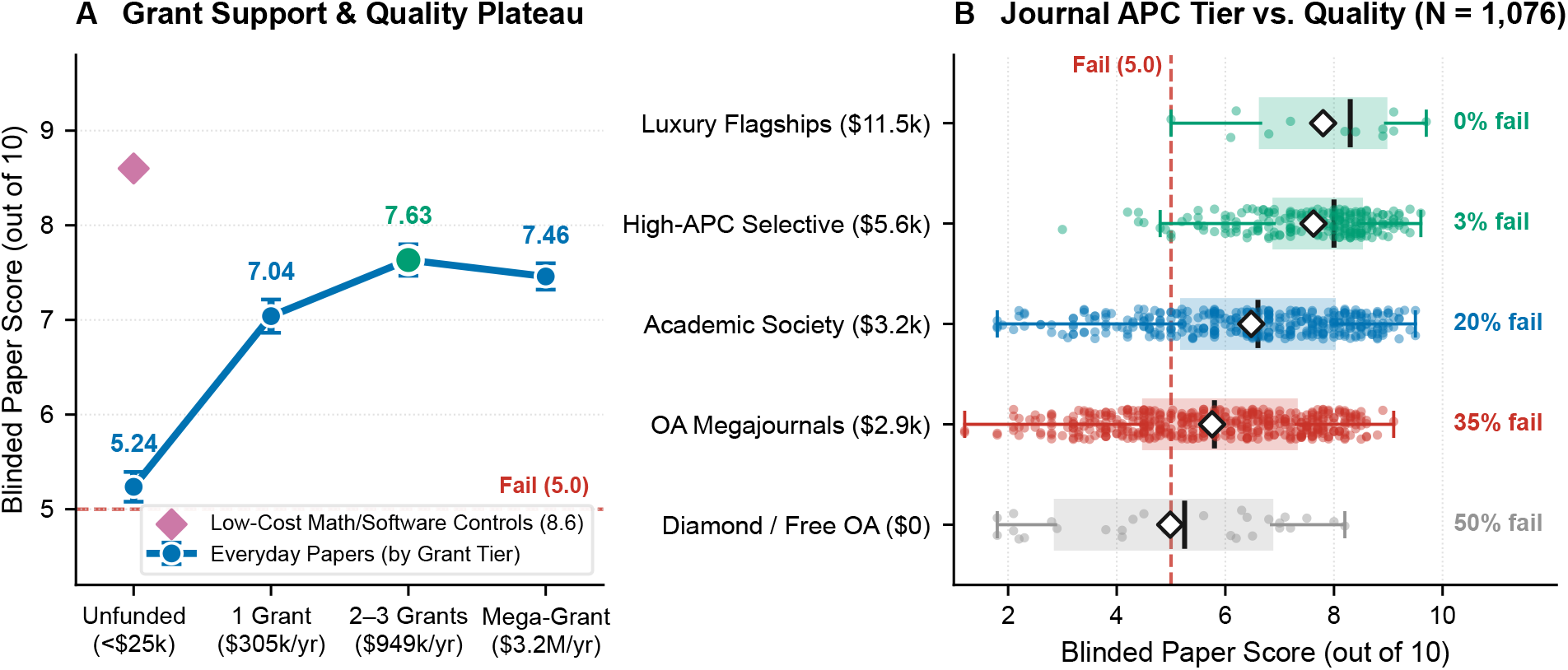
The economics of doing and publishing biomedical science across 1,076 everyday full-text papers. Panel A plots blinded BARS quality scores (/10) across unfunded studies and annual NIH grant tiers. Quality reaches an inflection point at 2–3 focused grants (median $949k/year, scoring 7.63*/*10, median 8.00) before plateauing in mega-grant consortia (*>* $1.5M/year, scoring 7.46*/*10). Low-cost mathematical and software Sound Controls (purple diamond) average 8.5*/*10 (0% failed), showing no BARS bias against inexpensive rigorous science. Panel B compares blinded scores across five journal Article Processing Charge (APC) tiers (*N* = 1, 076): commercial megajournals (*∼* $2, 880 APC) average 5.76*/*10 (35% failed), whereas similarly priced academic society journals (*∼* $3, 240 APC) average 6.48*/*10 (21% failed, *p* = 1.1 *×* 10*^−^*^8^).

First, BARS shows no detectable penalty against inexpensive science when methods and claims are aligned. Within our 50 Sound Controls (Cohort B), 14 low-cost mathematical, statistical, or software studies with zero wet-lab budget averaged 8.5 *±* 1.0 out of 10 (0% failed), matching the 36 expensive multi-assay wet-lab Sound Controls (8.7 *±* 0.8, *p* = 0.39, Figure 6A) and our lab’s lead-author theoretical papers (8.2*/*10). Conversely, BARS does not excuse flawed designs in heavily funded studies: the 18 NIH-funded Flawed Controls (Cohort A) carried a median grant portfolio of $2.07 million/year (mean $3.76 million/year) yet scored 5.3 out of 10 (4.4 across all 50 Flawed Controls).

In everyday empirical biology, however, study budget matters up to a clear inflection point because causal biological claims require experimental validation (Figure 6A, Supplementary Table S6). Across the 1,076 everyday papers, studies acknowledging zero external grants (*n* = 615) averaged 6.14 *±* 1.85 out of 10 (27.5% failed below 5.0; 6.12 *±* 1.83 and 27.9% failed across all *n* = 854 non-NIH papers, and 5.33 *±* 1.78 with 42.6% failed among *n* = 230 secondary database studies). Among the 222 NIH-funded everyday papers, single-grant studies (*<*$600k/year, median $305k/year, *n* = 80) rose to 7.04 *±* 1.57 (12.5% failed). Quality peaked at 7.63 *±* 1.45 (median 8.00, 9.2% failed) for focused teams supported by two to three grants totaling $0.6 to 1.5 million/year (median $949k/year, *n* = 76, *p* = 0.015 vs. single-grant studies). Beyond $0.95 million/year, additional funding yielded no gain in mean quality: mega-grant consortia (*>*$1.5 million/year, median $3.20 million/year, *n* = 66) plateaued at 7.46 *±* 1.13 (median 7.80, *p* = 0.42 vs. 2–3 grants, though outright failures below 5.0 dropped to 1.5%, 1*/*66, Fisher’s exact *p* = 0.013), and total acknowledged grant counts showed the same plateau (6.14 for 0 grants versus 7.35 for both 4–6 grants and *≥* 7 grants).

Comparing journal APCs against paper quality across all 1,076 everyday papers revealed a sharp business-model gap (Figure 6B, Supplementary Table S6).

- **Low scores at** $2, 900 **commercial megajournals.** Commercial APC-driven megajournals and open-threshold OA venues in the $1, 700–$3, 900 tier (*n* = 459, mean APC $2, 879, including *PLOS ONE*, *Scientific Reports*, *Frontiers*, *MDPI*, and *BMC* series) averaged 5.76 *±* 1.74 out of 10, with 35.1% failing below 5.0 (4.40 *±* 1.50, 64.9% failed in 2020–2025 Everyday Papers). Zero-fee diamond OA venues (*n* = 28) averaged 4.99 *±* 2.11 (50.0% failed).
- **Higher rigor per dollar at academic society journals.** Selective academic society and university press journals (*n* = 361, mean APC $3, 242, such as *Molecular Biology and Evolution*, *PNAS*, *PLOS Pathogens*, *PLOS Genetics*, *eLife*, and *Nucleic Acids Research*) charge nearly the same APC as commercial megajournals ($3, 242 vs. $2, 879) yet score +0.72 points higher (6.48 *±* 1.80, *p* = 1.1 *×* 10*^−^*^8^), cut failures to 20.5%, and deliver nearly 3*×* more peer-review improvement.
- **Diminishing returns at** $11, 500 **high-fee flagships.** High-APC selective journals ($4, 500–$7, 000, mean $5, 579, *n* = 216, such as *Nature Communications*, *Science Advances*, and *Genome Biology*) averaged 7.62 *±* 1.17 (3.2% failed). Top-fee flagships charging $9, 500–$12, 290 (mean $11, 528, *n* = 12 everyday, *n* = 51 overall, such as *Nature*, *Science*, and *Cell*) cost an extra $5, 950 per paper for a non-significant +0.18-point difference (7.80 *±* 1.49, *p* = 0.69), and among Sound Controls (Cohort B) $3, 230 society journals (8.74) slightly outscored $11, 738 flagship journals (8.65).

We next measured the token consumption and API cost of BARS across our reviews and compared them against human peer-review labor and the broader downstream costs of flawed publications (Supplementary Table S6).

- **Token and API cost to review a paper with BARS (4.1 to 8.7 cents) and extrapolation to all of PubMed.** A full-text BARS review of the main article text, equations, and inline tables (excluding external supplementary attachments) averages 22, 119 tokens (15, 392 input tokens for the manuscript and rubric plus 6, 727 reasoning and output tokens). At standard frontier API pricing, a single review costs $0.041 in Batch mode ($0.087 on-demand), a 3-run ensemble costs $0.124, and our 1,874-review study cost $77.48 in Batch mode ($162.10 on-demand). Auditing every open-access paper deposited in PubMed Central in 2025 (921, 565 articles) would cost $38,109 per year in Batch mode. Auditing the entire annual flow of PubMed (1.97 million articles, assuming full-text access) would cost $81,272 per year—roughly the publication fees of seven high-APC flagship papers—and auditing the entire 10.9-million-paper historical full-text archive of PubMed Central would cost $450,743 (Supplementary Table S6).
- **Human peer-review labor (15 hours and $1,505–$2,630 per paper vs. $0.04 BARS).** Across our 100 open-review papers, human review required 2.61 referees per paper (median 3, 213 referee words), consuming *∼* 15.05 hours of researcher and editor time. Using the conservative global benchmark of $100/hour from Aczel et al. [44] yields $1,505 per paper (29.6 million hours or $2.96 billion/year across PubMed). Using median US R1 biomedical faculty compensation (AAUP and AAMC median associate-to-full professor base salaries of $145k–$210k plus 30% fringe benefits, or *∼* $130–$175/hour of faculty time) raises the human review cost to $1,950–$2,630 per paper ($3.8–$5.2 billion/year across PubMed). A 4-cent BARS pre-screen takes under three minutes, is 36, 000*×* to 64, 000*×* cheaper than human referee labor, recalls 97.9% of unresolved human critiques, and catches 3.85 additional quantitative flaws per paper.
- **Downstream costs of preventable flaws vs. a 4-cent pre-screen.** Although acknowledged grants typically fund multi-paper research programs instead of single studies, publishing preventable methodological flaws still carries substantial direct and downstream costs. Across our 1,076 everyday papers alone, authors and funders spent $830, 000 directly on APCs for 256 failing studies scoring below 5.0. At a systemic level, irreproducible preclinical research is estimated to waste $28 billion per year in the United States [46, 47]. Even a single flawed or falsified landmark paper can misdirect an entire field—such as the retracted 2006 *Nature* A*β*\*56 Alzheimer’s oligomer study, which preceded over $287 million in follow-up NIH oligomer awards [48]—or require years of costly cancer-microbiome re-analyses [11, 12, 45]. Catching analytical and control flaws prior to publication with a 4-cent BARS pre-screen is orders of magnitude cheaper than correcting them after the fact.

## 3 Discussion

Scientific publishing currently lacks an automated quality floor. Across our full-text reviews, we showed that BARS—stripped of author names, institutions, journal titles, and dates—separates Sound Controls from retracted or refuted Flawed Controls with 97% to 99% accuracy, zero false rejections of Sound Controls at the 5.0-point threshold, and more than twice the test-retest consistency of human referees. Applying this calibrated yardstick across everyday biomedical papers published between 1991 and 2025 yielded six core lessons:

1. **Blinding matters:** Famous authors and flagship journals enjoy a +0.35-point prestige halo that disappears when metadata are stripped.
2. **Roughly half of recent papers fail on rigor:** Across 320 recent 2020–2025 papers spanning eight fields, 54.1% score below 5.0, led by unverified data-mining templates and contamination-prone assays.
3. **Peer review works at selective journals, not megajournals:** Across 50 preprint-to-published pairs, revisions at selective flagship and society journals improved papers more than twice as much as at megajournals (+0.38 vs. +0.18 Core points; +0.62 vs. +0.30*/*10 overall), and unreviewed `bioRxiv`/`medRxiv` preprints (6.46*/*10) outscored published megajournal articles (4.18–5.76*/*10). Across 100 open-review papers, BARS recalled 97.9% of unresolved human critiques while exposing referee anomalies in 58% of megajournal dossiers vs. 2% at selective venues.
4. **Ecosystem shifts drove the 35-year quality decline:** Most (54%–58%) of the 1.82-point drop between the 1990s (7.63) and 2020s (5.81) came from two waves of compositional shift—early-2000s candidate-gene/microarray studies and commercial OA series, followed by 2016–2025 megajournals and secondary database templates—whereas within-journal drift was small (*−*0.23 points/decade).
5. **Self-auditing works—including on our own papers and this manuscript:** Blinded review across 238 career papers by two senior authors (164 by S.L.K.P.; 74 by A.N.) caught real mathematical, control, and table errors in both our lead-author and middle-author studies. It also scored a blinded draft of this manuscript 7.8*/*10 and caught five statistical and consistency flaws that we fixed before publication.
6. **Perceived prestige diverges from actual scientific quality and value:** Mega-grant consortia (*>* $1.5M/year) and $11, 500 high-fee flagship APCs do not buy higher rigor than focused 2–3 grant labs (*∼* $0.95M/year, 7.63 vs. 7.46*/*10) or $3, 240 society journals (8.74 vs. 8.65*/*10 on Sound Controls). Society journals also outscore $2, 880 commercial megajournals by +0.72 points, and a 4-cent BARS pre-screen can audit all of PubMed for $81,272/year.

### Why Do Some Flawed Papers Get Highly Cited? Four Archetypes of Highly Cited Failures

Although higher-scoring papers earn substantially more citations on average (mean 176.3 citations for papers scoring *≥* 8.0 vs. 12.5 for papers scoring *<* 5.0; Supplementary Figure S1), citation count alone is a poor proxy for technical validity. Setting aside our 50 Flawed Controls (Cohort A), 8 verified full-text papers that failed below 5.0 still accumulated *≥* 100 citations and 19 accumulated *≥* 50 citations, alongside recent 2024–2025 failures already accruing 15–39 citations per year (Supplementary Figure S1A; browsable via the “Highly Cited Failures” filter at https://veg.github.io/BARS/). Almost all fall into four archetypes.

1. **First-Wave Crisis Gold-Rush Papers (**100**–**156 **citations).** Early pandemic papers making bold claims about viral origins or drug repurposing accumulate rapid citations before scrutiny catches up. Examples include `TIME_ANON_0224` (3.7*/*10, 156 citations, claiming SARS-CoV-2/MERS recombination from an unmasked 7-sequence alignment), `TIME_ANON_0922` (2.2*/*10, 139 citations, docking Lopinavir and Ritonavir simultaneously into the same protease site), and `SP_ANON_103` (2.2*/*10, 55 citations, mistaking PCR carryover for 2019 Italian COVID-19 cases) [17].
2. **Useful Gene-Family Catalogs With Broken Downstream Math (**73**–**158 **citations).** Many genome-era studies are cited for their Table 1 gene nomenclature despite flawed evolutionary statistics. Examples include `TIME_ANON_08` (4.8*/*10, 158 citations, plotting PAML’s `dN/dS = 99` divide-by-zero error code as literal positive selection *ω* = 99) and `TIME_ANON_0605` (4.9*/*10, 73 citations, which retained a copy-pasted “MAPK gene family” conclusion inside a millet *MYB* paper).
3. **Prestige-Halo Claims in Flagship Venues (**62**–**556 **citations).** Provocative claims in high-impact journals attract both uncritical citations and refutations. Examples include `SP_ANON_109` (*Nature* 2016, 4.1*/*10, 556 citations, claiming ongoing HIV-1 replication under therapy alongside a negative evolutionary rate table) and `TIME_ANON_0867` (4.5*/*10, 111 citations, claiming cisplatin induces promoter methylation in vivo when 0*/*18 treated genes became hypermethylated).
4. **Buzzword-Combo Templates Cited by Other Templates (**15**–**39 **citations/year).** Recent 2024–2025 papers chaining single-cell RNA-seq, Mendelian randomization, LASSO Cox signatures, and molecular docking (`C21`, 2.8*/*10, 39 citations; `C86`, 4.4*/*10, 36 citations) are rapidly cited as boilerplate introduction references by hundreds of other formulaic pipeline papers.

### Why These Four Metrics? Epistemic Invariance as Technologies Change

A central assumption of any literature audit is the choice of grading metrics: why evaluate papers on methodological rigor (3.5 points), claim-evidence alignment (2.5 points), biological plausibility (2.0 points), and reproducibility (2.0 points), and would our conclusions change if those metrics shifted in the future? We designed BARS around these four pillars instead of technology-specific checklists because laboratory assays and computational tools turn over every few years (from restriction enzymes and microarrays in the 1990s to bulk RNA-seq, single-cell transcriptomics, and deep-learning structure prediction today), whereas the core logic of scientific inference does not change. Regardless of the decade or assay, a valid empirical study must still (i) include appropriate controls and avoid statistical leakage or pseudoreplication, (ii) restrict its causal claims to what its data actually test, (iii) respect basic physical, chemical, and biological constraints, and (iv) describe its procedures and data clearly enough to be checked.

In practice, our empirical findings are highly robust to changes in rubric weights or threshold definitions for three reasons.

1. **Invariance to removing evolving open-science norms.** The one dimension whose community norms have genuinely shifted over time is open data and code deposition. Removing the 2.0-point Reproducibility score altogether leaves the 8.0-point Scientific Core and preserves the main conclusions of our study. Specifically, BARS still separates Sound Controls from Flawed Controls with AUC = 0.97, selective peer review still improves manuscripts more than twice as much as megajournal review (+0.38 vs. +0.18 Core points), and the 35-year secular trend shows an even steeper decline (*−*0.68 scaled Core points/decade, *p* = 8.7 *×* 10*^−^*^29^).
2. **Single dominant rigor axis across sub-scores.** Across all 1,777 catalog reviews, the three Scientific Core sub-scores correlate at *r* = 0.93 to 0.97 and load onto a single dominant principal component capturing 89.4% of total variance (Supplementary Section S3). Consequently, any reasonable reweighting of methods, claim alignment, and biological plausibility yields nearly identical paper rankings (*r >* 0.99).
3. **Stability across diagnostic thresholds.** One can define failure strictly at *<* 5.0 (0% false alarms on Sound Controls), moderately at *<* 6.0 (2% false alarms), or broadly at *<* 6.5 (4% false alarms). Across all three cutoffs, the ranking of scientific fields, the 2.5-to-3.0 point gap between selective journals and commercial megajournals, and the funding inflection at 2–3 NIH grants remain unchanged (Supplementary Tables S1–S6).

### Guardrails and a Four-Part Prescription for Scientific Publishing and Training

Our results and recent automated screening benchmarks [37, 38, 39, 40, 41, 43] point to six practical guardrails for AI review. First, de-identification, workspace isolation, and calibration against known controls are essential; otherwise models reward author and journal prestige [35, 36] or over-flag 25% to 29% of clean Sound Controls [43]. Second, text-only AI reviewers cannot detect manipulated western blots, spliced microscopy images, or hidden ancestry biases inside external consortium databases, and open-access XML sampling (`OPEN_ACCESS:y AND IN_EPMC:y`) under-represents hybrid or subscription society journals that deposit fewer open-access XML articles in Europe PMC (such as *Systematic Biology* or *Evolution*). Third, automated post-review verification checks must confirm that every accession or identifier cited by an AI review actually appears in the manuscript. We discovered this failure mode early in our method development: when an initial version of our text scrubber replaced real database accession numbers with the placeholder `[ACCESSION_REDACTED]`, BARS guessed a plausible accession number in 0.99% of early reviews (17*/*1, 719). Preserving non-identifying accession format strings and adding an automated verification gate eliminated 100% of these cases, and across all 385 BARS-only catches in our open-review cohort, 100% (385*/*385) were verified directly in the text (Supplementary Section S2). Fourth, rubrics and cutoffs must be calibrated to domain norms and re-anchored on Cohort A and B controls across independent AI model families whenever switching or upgrading models. Fifth, the AI model’s objective function and system prompt must be locked against reward drift and commercial incentive misalignment. In human publishing, commercial megajournals drifted toward superficial review because APC revenue scales with acceptance volume, while consumer chatbots are trained via reinforcement learning to be agreeable and sycophantic to users. If an AI reviewer is tuned to please authors or maximize publisher throughput, or if authors optimize text to game a static prompt (Goodhart’s Law), its diagnostic value will collapse. Any production AI review system must therefore forbid sycophancy in its system prompt and undergo continuous, public verification against a frozen benchmark of retracted/refuted and verified controls (‘Cohort A/B‘), maintaining 100% specificity at *<* 5.0 and AUC *≥* 0.95. Sixth, pre-publication auditing must protect manuscript confidentiality and intellectual property. Funders and journals rightly prohibit pasting unpublished drafts into consumer web chatbots that log prompts for training. By contrast, BARS strips identifying headers locally before any text leaves the user’s machine and can protect manuscript confidentiality in three ways. It can run through enterprise zero-data-retention cloud APIs (already used by journal submission and plagiarism systems) that contractually forbid model training and delete inputs upon completion. It can also run automatically on public preprint servers (`bioRxiv`, `medRxiv`) where authors have already shared their text, or entirely offline on institutional GPUs using open-weights reasoning models re-calibrated against our public Cohort A and B benchmark.

Subject to those guardrails, we recommend four practical steps for authors, journals, funders, and doctoral training programs.

- **For Authors (Pre-Submission Auditing, Co-Author Due Diligence, and Annual Career Self-Audits):** Run a 3-minute BARS audit before submission to catch table contradictions, unit errors, and overstated titles, and audit the entire draft when joining a large collaboration as a middle co-author instead of checking only one’s own methods paragraph. Beyond single papers, individual scientists and laboratories can run blinded BARS self-audits annually across their published portfolios (as in Figure 5) to track methodological rigor over time and provide verifiable quality benchmarks in grant, hiring, and promotion dossiers.
- **For Journals and Societies (Automated Table/Math Checks and Referee Quality Control):** Pair human referees with a blinded BARS pre-screen to verify internal table math, degrees of freedom, and revision-induced errors while automatically flagging boilerplate checklists and coercive self-citation demands in incoming human referee reports.
- **For Funders and Universities (Reward Rigor Over Volume):** Shift incentives away from raw paper counts and high-fee APCs toward focused 2–3 grant research teams publishing in rigorous academic society journals.
- **For Doctoral Training and Trainee Peer Review (Study Design and Active Review Apprenticeship):** Our failure taxonomy across 1,076 everyday papers shows that modern papers rarely fail because trainees cannot run software packages. Instead, papers fail because of experimental design and statistical identification errors, including missing negative controls, pseudoreplication, circular feature selection, and causal overclaiming from secondary databases. Graduate curricula in quantitative biology should therefore prioritize adversarial study design, negative controls, and statistical identifiability over routine execution of software templates. Equally important, BARS should be used to *strengthen*, not replace, how trainees learn to review manuscripts. In journal clubs and mentored co-reviewing, trainees should first draft their own independent critique of a blinded paper and then compare it against a BARS audit. Because BARS handles exhaustive table arithmetic, *p*-value consistency, and reporting checks, it frees trainees and mentors to focus on deeper conceptual questions: whether a biological hypothesis is worth testing, what experiment would decisively establish a mechanism, and where an automated main-text screen requires domain nuance. Used this way, BARS turns peer-review training from unguided trial-and-error into structured forensic apprenticeship.

## 4 Methods

### 4.1 Study Design and Paper Sampling

We conducted six coordinated experiments totaling 1,874 full-text reviews of 1,714 unique open-access biomedical articles from PubMed Central and Europe PubMed Central using BARS (Blinded AI Review System; Table 1). To prevent post-hoc metric tuning, the four-category, 10-point BARS scoring rubric and category weights (`REVIEW_TEMPLATE.md`, commit `01665db`) were defined and frozen prior to sampling any manuscripts. In Experiment 1 (200-Paper Calibration Set), we assembled 200 papers across five core fields: viral evolution, cancer and microbiome, statistical genetics, single-cell transcriptomics, and molecular evolution. The set comprised 50 retracted or refuted papers with documented flaws (Flawed Controls, Cohort A), 50 verified sound benchmark studies (Sound Controls, Cohort B), and 100 uncurated routine papers from 2020–2025 (Everyday Papers, Cohort C). We first ran a 75-paper pilot subset (25 each from Cohorts A, B, and C, evaluated in both unblinded and blinded modes) to lock our scoring thresholds (‘< 5.0‘ for fundamental flaws, ‘< 6.0‘ for major flaws). We then tested those locked cutoffs on the strictly held-out 125-paper subset of the Calibration Set (25 Flawed Controls, 25 Sound Controls, and 75 Everyday Papers).

In Experiment 2 (Repeatability), 30 papers from the Calibration Set (10 each from Cohorts A, B, and C) were reviewed blind three separate times under fresh anonymous IDs (90 reviews). In Experiment 3 (Everyday Papers Across Eight Fields, *N* = 320), we expanded our 2020–2025 audit to *n* = 40 papers per field across all eight fields. This included *N* = 200 papers across the five core biomedical fields (with an exact 20/20 split between oncology and host-associated microbiome studies) and *N* = 120 papers across three unseen fields (ecology and macroevolution, neuroscience and neuroimaging, and AI drug discovery). In Experiment 4 (Peer Review Value-Added and Open-Review Concordance), we audited 50 matched pairs of version-1 preprints and their final published articles (100 blinded texts) across selective flagship and society journals (*n* = 25 pairs) and open-access megajournals (*n* = 25 pairs). We paired this before-and-after test with a balanced cohort of 100 open-review articles published between 2020 and 2025 (`OPEN_ANON_001`–`OPEN_ANON_100`, 10 per subfield across our five core domains). Fifty articles came from selective flagship or society open-review venues (*eLife*, *PLOS Biology*, *PLOS Genetics*, *PLOS Pathogens*, and *PLOS Computational Biology*) and 50 came from open-review megajournals (*PLOS ONE*), each paired with its verbatim human referee reports (*≥* 2.5 KB, 261 distinct human reviewers and 363 substantive critiques total). Blinded Stage 1 BARS reviews were locked before Stage 2 concordance auditing against the human referee reports. In Experiment 5 (35-Year Secular Trend and Economics), we sampled 1,000 date-blinded papers across seven 5-year windows from 1991–1995 to 2021–2025 and four biomedical subfield tracks, and strictly excluded 85 archival 1990s/early-2000s records whose Europe PMC XML lacked main body text (*N* = 915 verified full-text articles, 226–233 per track). Across the 915-paper temporal cohort and the 1,076-paper everyday economics corpus (the 915 temporal papers plus 161 initial everyday papers, or 1,236 with the 160 field-expansion papers), we analyzed study design drivers, within-journal fixed effects, Oaxaca–Blinder decompositions, NIH RePORTER grant awards, and journal APCs. In Experiment 6 (Blinded Self-Audit), we audited all open-access papers in PubMed Central co-authored by two of our senior authors: 164 papers by S.L.K.P. (2004–2026; *n* = 42 Lead, *n* = 122 Middle) and 74 papers by A.N. (2000–2026; *n* = 28 Lead, *n* = 46 Middle). This totaled 238 blinded reviews across 222 unique papers, after which we subjected a de-identified draft of this manuscript (`MANUSCRIPT_META_01`) to the same BARS pipeline.

### 4.2 Blinding Pipeline, Scoring Rubric, and Statistical Summary

Our automated preprocessor extracts the main article text, equations, and inline tables from each manuscript’s XML or HTML file while stripping the `<ref-list>` bibliography and excluding external supplementary attachments and external code repository contents (which are not fetched or executed). In Experiments 4 and 6b (and our unified JATS parser), `<back>` and `<custom-meta>` Data Availability notes are extracted alongside, whereas in Experiments 5 and 6a only XML was parsed; the 8-point Scientific Core (Core_8_) evaluates rigor, claims, and biology independently of data-availability links across all cohorts. The scrubber removes the article title, author names, affiliations, correspondence lines, email addresses, journal name, volume/page numbers, inline author self-citations (`Surname et al.` replaced with `[AUTHOR et al.]`), web identifiers (DOIs, PMIDs, PMCIDs, repository handles), and any retraction notices. For the 100 open-review manuscripts, decision letters and reviewer sub-articles are stripped prior to Stage 1 review. For the 35-year trend and career self-audits, the scrubber also strips all 4-digit years and recognizable lab software names. Each scrubbed document is graded by an isolated instance of `gemini-4-argon` with web search disabled using our standardized template (`REVIEW_TEMPLATE.md` ‘v1.1‘).

The rubric grades each paper out of 10.0 points across four categories: Methodological Rigor and Statistical Validity (3.5 points), Claim-Evidence Alignment (2.5 points), Biological Plausibility (2.0 points), and Reproducibility and Data Integrity (2.0 points). Summing the first three categories gives the 8.0-point Scientific Core, which measures scientific quality independently of modern open-data links. Across all 1,777 catalog records, the three Scientific Core sub-scores correlate strongly (*r* = 0.93–0.97) while Reproducibility correlates more moderately (*r* = 0.76–0.80); principal component analysis identifies a primary scientific-rigor factor (PC1 = 89.4% of variance) plus a secondary reproducibility axis (PC2 = 7.7%; Supplementary Section S3). All statistical comparisons—including receiver operating characteristic curves, prevalence adjustments [33], test-retest intraclass correlation coefficients [34], paired preprint-to-published *t*-tests and Wilcoxon tests, within-journal fixed-effect regressions, and Oaxaca–Blinder decompositions—were run in Python 3.11. Complete mathematical equations, XML scrubber specifications, and full sensitivity tables (Tables S1–S7, Supplementary Figure S1) are provided in the Supplementary Information.

### 4.3 Data Availability and Ethics

Our investigation analyzed only publicly available, open-access scientific texts from PubMed Central and Europe PubMed Central. No human subjects or biological samples were involved. All 1,874 review dossiers, verbatim human referee reports, Stage 2 concordance matrices, scrubbed inputs, metadata tables, and analysis scripts are openly available under MIT and CC-BY-4.0 licenses at https://github.com/veg/BARS and interactively searchable at https://veg.github.io/BARS/. A dedicated BARS tool will be available at the Galaxy instance at https://usegalaxy.org.

### 4.4 AI Disclosure Statement

Large language models are both the subject of this research and a tool used to conduct it. All 1,874 blinded manuscript evaluations and Stage 2 human-concordance audits were generated by isolated instances of `gemini-4-argon` (`Gemini 3.1 Pro`) operating inside the Antigravity agentic harness under the human authors’ direct experimental design and rubric (`REVIEW_TEMPLATE.md` ‘v1.1‘). We also used `gemini-4-argon` inside the Antigravity coding environment to assist with writing Python data-processing, statistical analysis, and figure-plotting scripts, building the interactive web explorer, and drafting and editing portions of the manuscript text and tables. The human authors directed the study design, reviewed and edited the manuscript, and audited the analysis pipelines and representative review outputs, taking full responsibility for the integrity and conclusions of the work.

## References

[1] Ioannidis, John P. A. (2005). Why Most Published Research Findings Are False. PLoS Medicine, 2(8), e124. doi:10.1371/journal.pmed.0020124

[2] Salter, Susannah J, Cox, Michael J, Turek, Elena M, Calus, Szymon T, Cookson, William O, Moffatt, Miriam F, Turner, Paul, Parkhill, Julian, Loman, Nicholas J, Walker, Alan W (2014). Reagent and laboratory contamination can critically impact sequence-based microbiome analyses. BMC Biology, 12(1). doi:10.1186/s12915-014-0087-z

[3] de Goffau, Marcus C., Lager, Susanne, Sovio, Ulla, Gaccioli, Francesca, Cook, Emma, Peacock, Sharon J., Parkhill, Julian, Charnock-Jones, D. Stephen, Smith, Gordon C. S. (2019). Human placenta has no microbiome but can contain potential pathogens. Nature, 572(7769), 329–334. doi:10.1038/s41586-019-1451-5

[4] Kennedy, Katherine M., de Goffau, Marcus C., Perez-Muñoz, Maria Elisa, Arrieta, Marie-Claire, Bäckhed, Fredrik, Bork, Peer, Braun, Thorsten, Bushman, Frederic D., Dore, Joel, de Vos, Willem M., Earl, Ashlee M., Eisen, Jonathan A., Elovitz, Michal A., Ganal-Vonarburg, Stephanie C., Gänzle, Michael G., Garrett, Wendy S., Hall, Lindsay J., Hornef, Mathias W., Huttenhower, Curtis, Konnikova, Liza, Lebeer, Sarah, Macpherson, Andrew J., Massey, Ruth C., McHardy, Alice Carolyn, Koren, Omry, Lawley, Trevor D., Ley, Ruth E., O’Mahony, Liam, O’Toole, Paul W., Pamer, Eric G., Parkhill, Julian, Raes, Jeroen, Rattei, Thomas, Salonen, Anne, Segal, Eran, Segata, Nicola, Shanahan, Fergus, Sloboda, Deborah M., Smith, Gordon C. S., Sokol, Harry, Spector, Tim D., Surette, Michael G., Tannock, Gerald W., Walker, Alan W., Yassour, Moran, Walter, Jens (2023). Questioning the fetal microbiome illustrates pitfalls of low-biomass microbial studies. Nature, 613(7945), 639–649. doi:10.1038/s41586-022-05546-8

[5] Sohail, Mashaal, Maier, Robert M, Ganna, Andrea, Bloemendal, Alex, Martin, Alicia R, Turchin, Michael C, Chiang, Charleston WK, Hirschhorn, Joel, Daly, Mark J, Patterson, Nick, Neale, Benjamin, Mathieson, Iain, Reich, David, Sunyaev, Shamil R (2019). Polygenic adaptation on height is overestimated due to uncorrected stratification in genome-wide association studies. eLife, 8. doi:10.7554/elife.39702

[6] Berg, Jeremy J, Harpak, Arbel, Sinnott-Armstrong, Nasa, Joergensen, Anja Moltke, Mostafavi, Hakhamanesh, Field, Yair, Boyle, Evan August, Zhang, Xinjun, Racimo, Fernando, Pritchard, Jonathan K, Coop, Graham (2019). Reduced signal for polygenic adaptation of height in UK Biobank. eLife, 8. doi:10.7554/elife.39725

[7] Zaidi, Arslan A, Mathieson, Iain (2020). Demographic history mediates the effect of stratification on polygenic scores. eLife, 9. doi:10.7554/elife.61548

[8] Squair, Jordan W., Gautier, Matthieu, Kathe, Claudia, Anderson, Mark A., James, Nicholas D., Hutson, Thomas H., Hudelle, Rémi, Qaiser, Taha, Matson, Kaya J. E., Barraud, Quentin, Levine, Ariel J., La Manno, Gioele, Skinnider, Michael A., Courtine, Grégoire (2021). Confronting false discoveries in single-cell differential expression. Nature Communications, 12(1). doi:10.1038/s41467-021-25960-2

[9] Zimmerman, Kip D., Espeland, Mark A., Langefeld, Carl D. (2021). A practical solution to pseudoreplication bias in single-cell studies. Nature Communications, 12(1). doi:10.1038/s41467-021-21038-1

[10] Li, Yumei, Ge, Xinzhou, Peng, Fanglue, Li, Wei, Li, Jingyi Jessica (2022). Exaggerated false positives by popular differential expression methods when analyzing human population samples. Genome Biology, 23(1). doi:10.1186/s13059-022-02648-4

[11] Poore, Gregory D., Kopylova, Evguenia, Zhu, Qiyun, Carpenter, Carolina, Fraraccio, Serena, Wandro, Stephen, Kosciolek, Tomasz, Janssen, Stefan, Metcalf, Jessica, Song, Se Jin, Kanbar, Jad, Miller-Montgomery, Sandrine, Heaton, Robert, Mckay, Rana, Patel, Sandip Pravin, Swafford, Austin D., Knight, Rob (2020). RETRACTED ARTICLE: Microbiome analyses of blood and tissues suggest cancer diagnostic approach. Nature, 579(7800), 567–574. doi:10.1038/s41586-020-2095-1

[12] Gihawi, Abraham, Ge, Yuchen, Lu, Jennifer, Puiu, Daniela, Xu, Amanda, Cooper, Colin S., Brewer, Daniel S., Pertea, Mihaela, Salzberg, Steven L. (2023). Major data analysis errors invalidate cancer microbiome findings. mBio, 14(5). doi:10.1128/mbio.01607-23

[13] Wei, Xinzhu, Nielsen, Rasmus (2019). CCR5-Δ32 is deleterious in the homozygous state in humans. Nature Medicine, 25(6), 909–910. doi:10.1038/s41591-019-0459-6

[14] Maier, Robert, Akbari, Ali, Wei, Xinzhu, Patterson, Nick, Nielsen, Rasmus, Reich, David (2019). No statistical evidence for an effect of CCR5-Δ32 on lifespan in the UK Biobank cohort. Nature Medicine, 26(2), 178–180. doi:10.1038/s41591-019-0710-1

[15] Cicchetti, Domenic V. (1991). The reliability of peer review for manuscript and grant submissions: A cross-disciplinary investigation. Behavioral and Brain Sciences, 14(1), 119–135. doi:10.1017/s0140525x00065675

[16] Marsh, Herbert W., Jayasinghe, Upali W., Bond, Nigel W. (2008). Improving the peer-review process for grant applications: Reliability, validity, bias, and generalizability.. American Psychologist, 63(3), 160–168. doi:10.1037/0003-066x.63.3.160

[17] Amendola, Antonella, Canuti, Marta, Bianchi, Silvia, Kumar, Sudhir, Fappani, Clara, Gori, Maria, Colzani, Daniela, Kosakovsky Pond, Sergei L., Miura, Sayaka, Baggieri, Melissa, Marchi, Antonella, Borghi, Elisa, Zuccotti, Gianvincenzo, Raviglione, Mario C., Magurano, Fabio, Tanzi, Elisabetta (2022). Molecular evidence for SARS-CoV-2 in samples collected from patients with morbilliform eruptions since late 2019 in Lombardy, northern Italy. Environmental Research, 215, 113979. doi:10.1016/j.envres.2022.113979

[18] Tang, Xiaolu, Wu, Changcheng, Li, Xiang, Song, Yuhe, Yao, Xinmin, Wu, Xinkai, Duan, Yuange, Zhang, Hong, Wang, Yirong, Qian, Zhaohui, Cui, Jie, Lu, Jian (2020). On the origin and continuing evolution of SARS-CoV-2. National Science Review, 7(6), 1012–1023. doi:10.1093/nsr/nwaa036

[19] Forster, Peter, Forster, Lucy, Renfrew, Colin, Forster, Michael (2020). Phylogenetic network analysis of SARS-CoV-2 genomes. Proceedings of the National Academy of Sciences, 117(17), 9241–9243. doi:10.1073/pnas.2004999117

[20] Aykut, Berk, Pushalkar, Smruti, Chen, Ruonan, Li, Qianhao, Abengozar, Raquel, Kim, Jacqueline I., Shadaloey, Sorin A., Wu, Dongling, Preiss, Pamela, Verma, Narendra, Guo, Yuqi, Saxena, Anjana, Vardhan, Mridula, Diskin, Brian, Wang, Wei, Leinwand, Joshua, Kurz, Emma, Kochen Rossi, Juan A., Hundeyin, Mautin, Zambrinis, Constantinos, Li, Xin, Saxena, Deepak, Miller, George (2019). The fungal mycobiome promotes pancreatic oncogenesis via activation of MBL. Nature, 574(7777), 264–267. doi:10.1038/s41586-019-1608-2

[21] Narunsky-Haziza, Lian, Sepich-Poore, Gregory D., Livyatan, Ilana, Asraf, Omer, Martino, Cameron, Nejman, Deborah, Gavert, Nancy, Stajich, Jason E., Amit, Guy, González, Antonio, Wandro, Stephen, Perry, Gili, Ariel, Ruthie, Meltser, Arnon, Shaffer, Justin P., Zhu, Qiyun, Balint-Lahat, Nora, Barshack, Iris, Dadiani, Maya, Gal-Yam, Einav N., Patel, Sandip Pravin, Bashan, Amir, Swafford, Austin D., Pilpel, Yitzhak, Knight, Rob, Straussman, Ravid (2022). Pan-cancer analyses reveal cancer-type-specific fungal ecologies and bacteriome interactions. Cell, 185(20), 3789–3806.e17. doi:10.1016/j.cell.2022.09.005

[22] Sebastiani, Paola, Solovieff, Nadia, DeWan, Andrew T., Walsh, Kyle M., Puca, Annibale, Hartley, Stephen W., Melista, Efthymia, Andersen, Stacy, Dworkis, Daniel A., Wilk, Jemma B., Myers, Richard H., Steinberg, Martin H., Montano, Monty, Baldwin, Clinton T., Hoh, Josephine, Perls, Thomas T. (2012). Genetic Signatures of Exceptional Longevity in Humans. PLoS ONE, 7(1), e29848. doi:10.1371/journal.pone.0029848

[23] Turchin, Michael C, Chiang, Charleston WK, Palmer, Cameron D, Sankararaman, Sriram, Reich, David, Hirschhorn, Joel N (2012). Evidence of widespread selection on standing variation in Europe at height-associated SNPs. Nature Genetics, 44(9), 1015–1019. doi:10.1038/ng.2368

[24] Berg, Jeremy J., Coop, Graham (2014). A Population Genetic Signal of Polygenic Adaptation. PLoS Genetics, 10(8), e1004412. doi:10.1371/journal.pgen.1004412

[25] Starr, Tyler N., Greaney, Allison J., Hilton, Sarah K., Ellis, Daniel, Crawford, Katharine H.D., Dingens, Adam S., Navarro, Mary Jane, Bowen, John E., Tortorici, M. Alejandra, Walls, Alexandra C., King, Neil P., Veesler, David, Bloom, Jesse D. (2020). Deep Mutational Scanning of SARS-CoV-2 Receptor Binding Domain Reveals Constraints on Folding and ACE2 Binding. Cell, 182(5), 1295–1310.e20. doi:10.1016/j.cell.2020.08.012

[26] Murrell, Ben, Wertheim, Joel O., Moola, Sasha, Weighill, Thomas, Scheffler, Konrad, Kosakovsky Pond, Sergei L. (2012). Detecting Individual Sites Subject to Episodic Diversifying Selection. PLoS Genetics, 8(7), e1002764. doi:10.1371/journal.pgen.1002764

[27] Murrell, Ben, Weaver, Steven, Smith, Martin D., Wertheim, Joel O., Murrell, Sasha, Aylward, Anthony, Eren, Kemal, Pollner, Tristan, Martin, Darren P., Smith, Davey M., Scheffler, Konrad, Kosakovsky Pond, Sergei L. (2015). Gene-Wide Identification of Episodic Selection. Molecular Biology and Evolution, 32(5), 1365–1371. doi:10.1093/molbev/msv035

[28] Jones, Felicity C., Grabherr, Manfred G., Chan, Yingguang Frank, Russell, Pamela, Mauceli, Evan, Johnson, Jeremy, Swofford, Ross, Pirun, Mono, Zody, MC, White, Simon, Birney, Ewan, Searle, Stephen, Schueler, Markus, Clamp, Michele, Di Palma, Federica, Lindblad-Toh, Kerstin, Kingsley, David M. (2012). The genomic basis of adaptive evolution in threespine sticklebacks. Nature, 484(7392), 55–61. doi:10.1038/nature10944

[29] Love, Michael I, Huber, Wolfgang, Anders, Simon (2014). Moderated estimation of fold change and dispersion for RNA-seq data with DESeq2. Genome Biology, 15(12). doi:10.1186/s13059-014-0550-8

[30] Field, Yair, Boyle, Evan A, Telis, Natalie, Gao, Ziyue, Gaulton, Kyle J., Golan, David, Yengo, Loic, Rocheleau, Ghislain, Froguel, Philippe, McCarthy, Mark I., Pritchard, Jonathan K. (2016). Detection of human adaptation during the past 2000 years. Science, 354(6313), 760–764. doi:10.1126/science.aag0776

[31] Nejman, Deborah, Livyatan, Ilana, Fuks, Garold, Gavert, Nancy, Zwang, Yaara, Geller, Leore T., Rotter-Maskowitz, Aviva, Weiser, Roi, Mallel, Giuseppe, Gigi, Elinor, Meltser, Arnon, Douglas, Gavin M., Kamer, Iris, Gopalakrishnan, Vancheswaran, Dadosh, Tali, Levin-Zaidman, Smadar, Avnet, Sofia, Atlan, Tehila, Cooper, Zachary A., Arora, Reetakshi, Cogdill, Alexandria P., Khan, Md Abdul Wadud, Ologun, Gabriel, Bussi, Yuval, Weinberger, Adina, Lotan-Pompan, Maya, Golani, Ofra, Perry, Gili, Rokah, Merav, Bahar-Shany, Keren, Rozeman, Elisa A., Blank, Christian U., Ronai, Anat, Shaoul, Ron, Amit, Amnon, Dorfman, Tatiana, Kremer, Ran, Cohen, Zvi R., Harnof, Sagi, Siegal, Tali, Yehuda-Shnaidman, Einav, Gal-Yam, Einav Nili, Shapira, Hagit, Baldini, Nicola, Langille, Morgan G. I., Ben-Nun, Alon, Kaufman, Bella, Nissan, Aviram, Golan, Talia, Dadiani, Maya, Levanon, Keren, Bar, Jair, Yust-Katz, Shlomit, Barshack, Iris, Peeper, Daniel S., Raz, Dan J., Segal, Eran, Wargo, Jennifer A., Sandbank, Judith, Shental, Noam, Straussman, Ravid (2020). The human tumor microbiome is composed of tumor type–specific intracellular bacteria. Science, 368(6494), 973–980. doi:10.1126/science.aay9189

[32] Rabosky, Daniel L. (2014). Automatic Detection of Key Innovations, Rate Shifts, and Diversity-Dependence on Phylogenetic Trees. PLoS ONE, 9(2), e89543. doi:10.1371/journal.pone.0089543

[33] Rogan, Walter J., Gladen, Beth (1978). ESTIMATING PREVALENCE FROM THE RESULTS OF A SCREENING TEST. American Journal of Epidemiology, 107(1), 71–76. doi:10.1093/oxfordjournals.aje.a112510

[34] Shrout, Patrick E., Fleiss, Joseph L. (1979). Intraclass correlations: Uses in assessing rater reliability.. Psychological Bulletin, 86(2), 420–428. doi:10.1037/0033-2909.86.2.420

[35] Tomkins, Andrew, Zhang, Min, Heavlin, William D. (2017). Reviewer bias in single-versus double-blind peer review. Proceedings of the National Academy of Sciences, 114(48), 12708–12713. doi:10.1073/pnas.1707323114

[36] Huber, Jürgen, Inoua, Sabiou, Kerschbamer, Rudolf, König-Kersting, Christian, Palan, Stefan, Smith, Vernon L. (2022). Nobel and novice: Author prominence affects peer review. Proceedings of the National Academy of Sciences, 119(41). doi:10.1073/pnas.2205779119

[37] Nuijten, Michèle B., Hartgerink, Chris H. J., van Assen, Marcel A. L. M., Epskamp, Sacha, Wicherts, Jelte M. (2016). The prevalence of statistical reporting errors in psychology (1985–2013). Behavior Research Methods, 48(4), 1205–1226. doi:10.3758/s13428-015-0664-2

[38] Brown, Nicholas J. L., Heathers, James A. J. (2017). The GRIM Test. Social Psychological and Personality Science, 8(4), 363–369. doi:10.1177/1948550616673876

[39] Weissgerber, Tracey, Riedel, Nico, Kilicoglu, Halil, Labbé, Cyril, Eckmann, Peter, ter Riet, Gerben, Byrne, Jennifer, Cabanac, Guillaume, Capes-Davis, Amanda, Favier, Bertrand, Saladi, Shyam, Grabitz, Peter, Bannach-Brown, Alexandra, Schulz, Robert, McCann, Sarah, Bernard, Rene, Bandrowski, Anita (2021). Automated screening of COVID-19 preprints: can we help authors to improve transparency and reproducibility?. Nature Medicine, 27(1), 6–7. doi:10.1038/s41591-020-01203-7

[40] Checco, Alessandro, Bracciale, Lorenzo, Loreti, Pierpaolo, Pinfield, Stephen, Bianchi, Giuseppe (2021). AI-assisted peer review. Humanities and Social Sciences Communications, 8(1). doi:10.1057/s41599-020-00703-8

[41] Liang, Weixin, Zhang, Yuhui, Cao, Hancheng, Wang, Binglu, Ding, Daisy Yi, Yang, Xinyu, Vodrahalli, Kailas, He, Siyu, Smith, Daniel Scott, Yin, Yian, McFarland, Daniel A., Zou, James (2024). Can Large Language Models Provide Useful Feedback on Research Papers? A Large-Scale Empirical Analysis. NEJM AI, 1(8). doi:10.1056/aioa2400196

[42] Schroter, Sara, Black, Nick, Evans, Stephen, Godlee, Fiona, Osorio, Lyda, Smith, Richard (2008). What errors do peer reviewers detect, and does training improve their ability to detect them?. Journal of the Royal Society of Medicine, 101(10), 507–514. doi:10.1258/jrsm.2008.080062

[43] Brian, Dominikus, Pereira Hernández, Xavier Isidro, Yin, Hao, Koury, Jeffrey, Rust, Ruslan (2026). Large language models detect methodological errors that human peer review misses in clinical trials. medRxiv. doi:10.64898/2026.10.25.26366455

[44] Aczel, Balazs, Szaszi, Barnabas, Holcombe, Alex O. (2021). A billion-dollar donation: estimating the cost of researchers’ time spent on peer review. Research Integrity and Peer Review, 6(1), 14. doi:10.1186/s41073-021-00118-2

[45] Stern, Andrew M., Casadevall, Arturo, Steen, R. Grant, Fang, Ferric C. (2014). Financial costs and personal consequences of research misconduct resulting in retracted publications. eLife, 3, e02956. doi:10.7554/eLife.02956

[46] Freedman, Leonard P., Cockburn, Iain M., Simcoe, Timothy S. (2015). The Economics of Reproducibility in Preclinical Research. PLOS Biology, 13(6), e1002165. doi:10.1371/journal.pbio.1002165

[47] Begley, C. Glenn, Ellis, Lee M. (2012). Drug development: Raise standards for preclinical cancer research. Nature, 483(7391), 531–533. doi:10.1038/483531a

[48] Piller, Charles (2022). Blots on a field? Science, 377(6604), 358–363. doi:10.1126/science.add9993

