## Supplementary Information for "Auditing Biomedical Literature with Blinded AI Peer Review: Where Peer Review Works, Where It Fails, and How to Strengthen It"

#### Supplementary Methods and Technical Specifications

##### S1. Corpus Construction and Locus Definitions

For Experiment 1 (Calibration, $N = 200$), we constructed three cohorts across five biomedical subfields: Viral Evolution, Cancer and Low-Biomass Microbiome, Statistical Genetics, Transcriptomics and Computational Biology, and Molecular Evolution. Cohort A ($N = 50$, 10 per subfield) comprises open-access manuscripts in PubMed Central that were either formally retracted or refuted by peer-reviewed re-analyses demonstrating fatal analytical, contamination, or statistical defects. Each Cohort A entry in `calibration/cohorts_200.json` records the retraction notice Digital Object Identifier (DOI) or refutation citation and the defect category. It also records three boolean locus indicators distinguishing internal manuscript flaws from upstream consortium data artifacts.

- `defect_locus_giant_strict` ($N = 47$): Excludes the three height polygenic adaptation studies (`A11`, `A23`, `A24`). Their false-positive selection signals originated in geographic stratification inside upstream GIANT summary statistics.
- `defect_locus_pilot5` ($N = 45$): Excludes the five papers (`A07`, `A11`, `A19`, `A24`, `A25`). Their invalidation depended on external database or simulation artifacts invisible in the manuscript text.
- `defect_locus_broad6` ($N = 44$): Excludes the union of both sets (`A07`, `A11`, `A19`, `A23`, `A24`, `A25`).

Cohort B ($N = 50$, 10 per subfield) comprises verified benchmark manuscripts selected for validated experimental design, mathematical rigor, or definitive forensic correction of prior field-wide errors, balanced to include at least 80% empirical biological discovery or forensic validation studies (43/50) and at most two foundational methods papers per subfield (7/50). Cohort C ($N = 100$, 20 per subfield) was sampled from Europe PubMed Central open-access full-text research articles published between 2020 and 2025 (`OPEN_ACCESS:y AND PUB_YEAR:[2020 TO 2025]`). Because the Europe PMC query requires open-access XML deposits (`OPEN_ACCESS:y AND IN_EPMC:y`), subscription or hybrid academic society journals that deposit fewer open-access XML articles in Europe PMC (such as *Systematic Biology*, *Evolution*, and *Molecular Ecology*) have lower sampling representation in automated Europe PMC draws than gold open-access venues. Phase 1 (`ANON_01` to `ANON_75`) served as the 75-paper pilot (25 per cohort), and Phase 2 (`ANON_76` to `ANON_200`) served as the strictly held-out 125-paper expansion (25 Cohort A, 25 Cohort B, 75 Cohort C). Across our multi-experiment catalog ($1,777$ records), three distinct megajournal subset counts appear in different analyses. Specifically, $n = 422$ is the number of strict commercial open-access megajournal papers (*PLOS ONE*, *Scientific Reports*, *Frontiers*, *MDPI*, *PeerJ*, and *Medicine*) inside the $N = 1,076$ everyday corpus (Table S4). Meanwhile, $n = 459$ is the number of papers in the $1.7k–$3.9k commercial OA APC price bracket inside the same $N = 1,076$ corpus (422 strict megajournals plus 37 field-specific commercial OA journals in that APC tier; Table S6). Finally, $n = 545$ is the total number of megajournal records across the entire multi-experiment catalog, which also includes the 50 megajournal papers in `open_review_exp`, 25 in `peer_review_exp`, and megajournal papers in the career self-audits and Cohort A. Across all six experiments, the $1,714$ unique papers map to $1,777$ catalog records (accounting for 29 cross-cohort natural replicates, 16 joint self-audit papers evaluated in both author batches, and 18 calibration/self-audit overlaps) and $1,874$ total full-text reviews (which additionally include 60 test-retest repeatability replicates, 25 Phase-1 unblinded pilot runs, 6 clinical-trial benchmark runs, and 6 meta-audit/ad-hoc evaluations).

#### S2. Deterministic XML Ingestion and De-Identification Scrubber

Full-text manuscripts were ingested from NCBI PubMed Central and Europe PubMed Central JATS XML and HTML endpoints via `scripts/fetch_papers.py` and `scripts/build_deidentified.py`. External supplementary binary attachments and external code repository contents are not fetched or executed; the Reproducibility sub-score evaluates stated data/code availability and internal numerical completeness, while $\text{Core}_8$ evaluates scientific rigor independently of availability links. To prevent inline mathematical parameters from dropping during XML parsing, our parser extracts both `<tex-math>` and `<mml:math>` nodes inside `<inline-formula>` and `<disp-formula>` elements. In Experiments 4 and 6b (and the unified JATS extractor in `scripts/build_deidentified.py`), `<back>` and front-matter `<custom-meta>` Data/Code Availability sections are extracted alongside `<body>` (preserving anchor `href` URLs prior to handle scrubbing), whereas in Experiments 5 and 6a only `<body>` XML was parsed. The deterministic scrubber strips:

1. Article titles, running headers, journal names, volume and issue strings, and copyright footers.
2. All author names, ORCID identifiers, email addresses (`[EMAIL_REDACTED]`), correspondence lines (`[CORRESPONDENCE_REDACTED]`), institutional affiliations, funding grants, Acknowledgements or Author Contributions sections, the `<ref-list>` bibliography (References section), and inline author self-citations in the body text (`<Surname> et al.` replaced with `[AUTHOR et al.]` and distinctive co-author surnames replaced with `[AUTHOR]`).
3. All Digital Object Identifiers, PubMed IDs, PMCIDs, preprint identifiers, and GitHub, GitLab, Bitbucket, UseGalaxy, or WorkflowHub URLs carrying user or lab handles (`[CODE_REPO_REDACTED]`, `[GALAXY_URL_REDACTED]`).

4. Explicit retraction prefixes (`RETRACTED:`, `WITHDRAWN:`) and inline publisher retraction notices.

5. In temporal-blinding runs (Experiment 5) and self-audit runs (Experiment 6), all 4-digit publication, copyright, or inline citation years (replaced with `[YEAR]`) and self-identifying laboratory software names (`HyPhy`, `Datamonkey`, `HIV-TRACE`, `Galaxy`, `UseGalaxy`, `Planemo`, `DuNovo` replaced with `[SOFTWARE]`). In initial review waves prior to commit `a61d8eb`, third-party `github.com/<org>/<repo>` links in 75 of 1,874 blinded reviews (4.0%) were redacted in post-processing rather than pre-processing. The file `blinding_url_sensitivity.json` lists all 75 review IDs and confirms that excluding them changes every cohort mean by at most 0.15 points (4.42 vs. 4.37 in Cohort A, 8.64 vs. 8.64 in Cohort B, 6.57 vs. 6.56 in the 35-year cohort, and 7.53 vs. 7.45 in the S.L.K.P. self-audit).

To verify that BARS critiques and accession citations are grounded in the manuscript text rather than hallucinated from training memory, we implemented an automated fact-verification script (`scripts/verify_review_facts.py`, integrated into continuous integration via `.github/workflows/check-ledger.yml` and archived in `fact_verification_ledger.json`). Across all 385 `ai_only_critical_catches` in the 100-paper open-review cohort, 385/385 (100.0%) were verified against the scrubbed manuscript text, and 115/115 (100.0%) of accession or rsID identifiers cited in those reviews matched the source manuscript verbatim. Across all $1,974$ accession or rsID tokens cited in all $1,719$ active full-text reviews, $1,944/1,974$ (98.48%) matched the source manuscript or were explicitly cited as external benchmark comparisons. Only 30 tokens across 17 reviews (0.99% of reviews) represented guessed accessions in Reproducibility bullets where an earlier scrubber had replaced the manuscript's accession with `[ACCESSION_REDACTED]`. All 17 cases were corrected to cite `[ACCESSION_REDACTED in scrubbed input]` or the exact manuscript accession (100.0% post-correction concordance). SHA-256 digests of all scrubbed input files are archived in `repeatability/scrubbed_inputs_sha256.json` and `ood/scrubbed_inputs_sha256.json`.

#### S3. Psychometric Scoring Equations and Statistical Estimators

Each scrubbed manuscript was evaluated by an isolated instance of `gemini-4-argon` invoked via stateless subagent contexts ('1' manuscript per context window, 'search_web' disabled) governed by `REVIEW_TEMPLATE.md` ('v1.1'). The quantitative scorecard sums four dimensions to 10.0 points:

$$S_{\text{total}} = S_{\text{rigor}} \in [0, 3.5] + S_{\text{claims}} \in [0, 2.5] + S_{\text{biology}} \in [0, 2.0] + S_{\text{repro}} \in [0, 2.0], \tag{S1}$$

alongside the 8.0-point Reproducibility-free Scientific Core:

$$\text{Core}_8 = S_{\text{rigor}} + S_{\text{claims}} + S_{\text{biology}} \in [0, 8.0]. \tag{S2}$$

Across the 200-paper calibration dataset, unweighted equal-dimension summation correlates with $S_{\text{total}}$ at $r = 0.9986$ and alters ROC-AUC by less than 0.0004. Across all $1,777$ catalog records ($1,703$ prior to the 74-paper A.N. self-audit expansion), pairwise Pearson correlations among the four sub-scores are $r(\text{Rigor}, \text{Claims}) = 0.971$, $r(\text{Rigor}, \text{Biology}) = 0.926$, $r(\text{Claims}, \text{Biology}) = 0.927$, $r(\text{Rigor}, \text{Repro}) = 0.800$, $r(\text{Claims}, \text{Repro}) = 0.785$, and $r(\text{Biology}, \text{Repro}) = 0.755$. Principal component analysis of the correlation matrix yields variance proportions PC1 = 89.43% (88.96% on the initial $1,483$ records), PC2 = 7.72% (8.17%), PC3 = 2.19%, and PC4 = 0.65%, showing that the rubric captures one primary scientific-rigor factor ($\text{Core}_8$) plus a secondary reproducibility and data-availability axis (PC2).

True defect prevalence $\hat{\pi}_{\mathrm{RG}}$ in wild literature was estimated from observed failure prevalence $p_{\mathrm{obs}}$ using the Rogan-Gladen estimator:

$$\hat{\pi}_{\mathrm{RG}} = \mathrm{clip}_{[0,1]}\left(\frac{p_{\mathrm{obs}} + \widehat{\mathrm{Sp}} - 1}{\widehat{\mathrm{Se}} + \widehat{\mathrm{Sp}} - 1}\right). \tag{S3}$$

Test-retest repeatability across $N$ manuscripts and $K = 3$ waves was quantified via Shrout-Fleiss two-way random-effects absolute-agreement intraclass correlation:

$$\mathrm{ICC}(2,1) = \frac{\mathrm{MS}_R - \mathrm{MS}_E}{\mathrm{MS}_R + (K-1)\mathrm{MS}_E + \frac{K}{N}(\mathrm{MS}_C - \mathrm{MS}_E)}, \quad \mathrm{ICC}(2,K) = \frac{K \cdot \mathrm{ICC}(2,1)}{1 + (K-1)\mathrm{ICC}(2,1)}, \tag{S4}$$

where $\mathrm{MS}_R$, $\mathrm{MS}_C$, and $\mathrm{MS}_E$ denote between-paper, between-wave, and residual mean squares, respectively. All metrics are deterministically reproducible via `scripts/verify_repeatability_and_calibration.py`.

#### S4. Replication and Blinded Benchmark on Error-Seeded Clinical Trials (Brian et al. 2026)

To cross-validate our blinded review architecture against an independent external benchmark, we replicated the computational pipeline and open dataset of Brian, Pereira Hernández, Yin, Koury, and Rust (2026) (`medRxiv 2026.10.25.26366455v2`). Their study evaluated 15 LLMs on historical British Medical Journal test papers from Schroter et al. (2008) ($N = 180$ reviews) and 58 randomized clinical trials published in 2026 ($N = 1,391$ reviews across 14 planted error categories: 9 major [`M1-M9`] and 5 minor [`m1-m5`]). Executing their validation scripts (`reproduce.py`, `validate_submission.py`, and `validate_further_trials.py`) reproduced all saved-data tables and statistics with zero discrepancies. At the same time, our audit revealed four critical properties of uncalibrated open-ended LLM peer review.

1. **Severe sensitivity–specificity tradeoff across models:** Across the 15 models tested by Brian et al., sensitivity to planted errors correlated strongly and negatively with specificity on unmodified control manuscripts (Spearman $\rho = -0.771, p = 0.00076$ on the primary 2026 trials; $\rho = -0.757, p = 0.0011$ across all 58 trials). Flagship models with the highest sensitivity (`Opus 5.5`, `Fable 5`, `GPT-6.1 Sol`) falsely flagged 25.2% to 28.6% of the 14 error categories on unmodified control trials (specificity 71.4%–74.8%), most frequently accusing clean trials of unjustified conclusions (`M7`, 576 control flags), missing screen-failure reasons (`m2`, 447 flags), and missing Hawthorne-effect discussions (`m5`, 189 flags).

2. **Sample-size asymmetry and workspace context leakage in top-ranked models:** Although plotted at the top of the 58-trial comparison in Brian et al.'s Figure 2, `Opus 5.5`, `Fable 5`, and `Opus 5` were evaluated on only $n = 3$ trials in 2026 (`N`, `I`, `M`) and omitted from all 55 expanded trials (`T01-T60`). Moreover, the repository's forensic audit (`reviewer_tool_and_context_audit.json`) discloses that during those initial 3-trial runs, the agent harness loaded a workspace `MEMORY.md` file describing the "Schroter-2008 replication," resulting in 33.3% (16/48) of Experiment 1 and 37.5% (18/48) of Experiment 2 reasoning traces explicitly referencing "schroter" or "planted" errors.

3. **Checklist memorization in training weights:** In isolated memory probes (`SupplFigure3_values.csv`), `Opus 5.5` recalled 11 of the 14 official Schroter error types without seeing a manuscript (`Fable 5` and `Opus 5` recalled 7/14).

4. **Omission of a frontier Gemini Pro model:** While Anthropic and OpenAI were evaluated with flagship reasoning models spending up to 16.3 min and $110,200$ output tokens ($2.24/review), Google was represented solely by the lightweight fast model `Gemini 3.8 Flash` (1.4 min, $10,100$ tokens, $0.045/review).

To fill this gap under strict workspace isolation, we reconstructed the clean control and error-seeded versions of the first three expanded 2026 *Nature Medicine* trials from PubMed Central JATS XML (`T01` = `PMC13668674`, `T02` = `PMC13577891`, `T03` = `PMC13668648`). We then evaluated all six blinded manuscripts with `gemini-4-argon` (`Gemini 3.1 Pro`) using Brian et al.'s reviewer prompt plus our calibrated 10-point score block (Table S7).

### Supplementary Figures and Tables

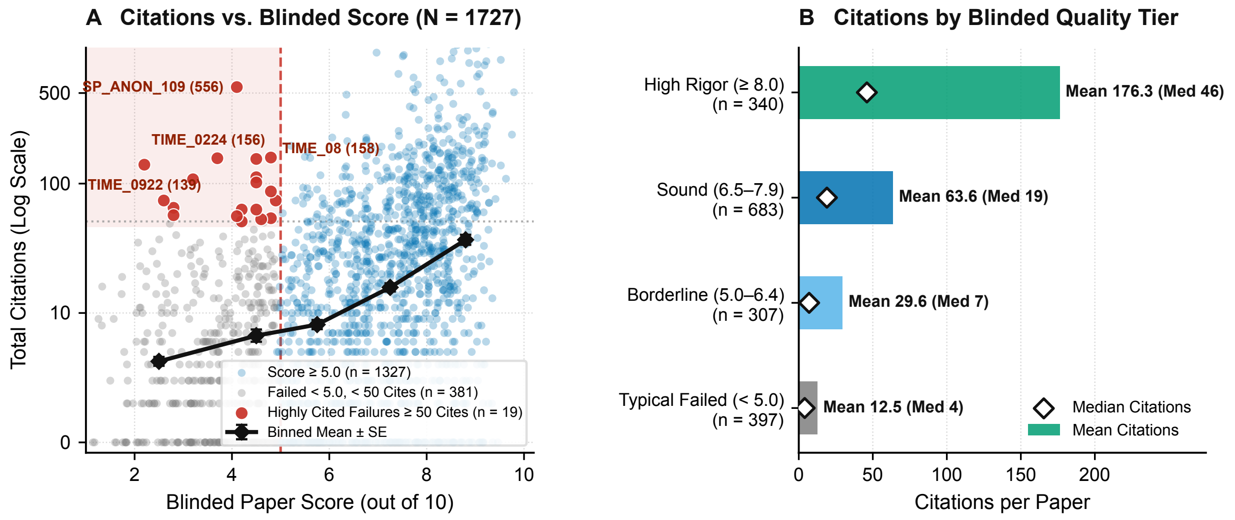


Figure S1: Relationship between blinded BARS quality scores and total citations across $N = 1,727$ non-retracted full-text catalog review records ($1,777$ catalog records minus 50 Cohort A Flawed Controls). Panel A plots $\log_{10}(\text{Citations} + 1)$ against the 10-point BARS score ($r = 0.30, p < 10^{-35}$, green binned mean $\pm$ SE curve). While higher-scoring articles attract substantially more citations on average, 19 articles that failed below the 5.0 rigor cutoff still accumulated $\geq 50$ citations (red shaded region, "Highly Cited Failures," labeled by their four archetypes in the main text). Panel B compares mean and median citations across four BARS quality tiers. High Rigor ($\geq 8.0$, $n = 455$) averages 176.3 citations (median 46) and Sound (6.5–7.9, $n = 589$) averages 63.6 (median 19), whereas Borderline (5.0–6.4, $n = 307$) averages 29.6 (median 7) and Typical Failed ($< 5.0$, $n = 376$) averages 12.5 (median 4).

Table S1: Complete calibration sensitivity matrix across Phase-1 pilot (`ANON_01-75`), strictly held-out expansion (`A26-A50, B26-B50`), upstream consortium exclusions, `PMC8135569` relabeling, and Cohort B control-composition subsets.

| Analysis Stratum | $N_A, N_B$ | Mean A | Mean B | ROC-AUC | $\widehat{\text{Se}}_{<5.0}$ | $\widehat{\text{Sp}}_{<5.0}$ | $\widehat{\text{Se}}_{<6.0}$ | $\widehat{\text{Sp}}_{<6.0}$ |
|---|---|---|---|---|---|---|---|---|
| Full 200-Paper Study (Published) | 50, 50 | $4.42 \pm 1.97$ | $8.61 \pm 0.93$ | 0.9662 | 66.0% | 100.0% | 76.0% | 98.0% |
| Phase-1 Pilot Only (`ANON_01-75`) | 25, 25 | $4.26 \pm 2.17$ | $8.20 \pm 1.41$ | 0.9232 | 68.0% | 96.0% | 76.0% | 92.0% |
| Held-Out Expansion (`A26-A50, B26-B50`) | 25, 25 | $4.59 \pm 1.78$ | $8.78 \pm 0.85$ | 0.9824 | 64.0% | 100.0% | 76.0% | 100.0% |
| Strict GIANT Exclusion (Excl. A11, 23, 24) | 47, 50 | $4.18 \pm 1.77$ | $8.61 \pm 0.93$ | 0.9832 | 70.2% | 100.0% | 80.9% | 98.0% |
| Internal Flaws Only (Excl. 5 Upstream) | 45, 50 | $3.99 \pm 1.54$ | $8.61 \pm 0.93$ | 0.9902 | 75.6% | 100.0% | 88.9% | 98.0% |
| Broad Upstream Exclusion (Excl. 6 Upstream) | 44, 50 | $3.91 \pm 1.46$ | $8.61 \pm 0.93$ | 0.9936 | 77.3% | 100.0% | 90.9% | 98.0% |
| Original Pilot Label (`PMC8135569` in B) | 49, 51 | $4.45 \pm 1.99$ | $8.51 \pm 1.18$ | 0.9536 | 65.3% | 98.0% | 75.5% | 96.1% |
| Excl. 9 Refutation Controls in B | 50, 41 | $4.42 \pm 1.97$ | $8.51 \pm 0.97$ | 0.9605 | 66.0% | 100.0% | 76.0% | 97.6% |
| Excl. 2 Senior-Author Controls in B | 50, 48 | $4.42 \pm 1.97$ | $8.60 \pm 0.94$ | 0.9654 | 66.0% | 100.0% | 76.0% | 97.9% |
| Independent Primary Controls Only in B | 50, 39 | $4.42 \pm 1.97$ | $8.49 \pm 0.99$ | 0.9592 | 66.0% | 100.0% | 76.0% | 97.4% |
| Specialist Society Controls Only in B | 50, 15 | $4.42 \pm 1.97$ | $8.42 \pm 0.88$ | 0.9640 | 66.0% | 100.0% | 76.0% | 100.0% |

Table S2: Per-cohort and pooled test-retest repeatability metrics ($N = 30 \times K = 3 = 90$ blinded evaluations). $SD_e$ denotes the two-way residual error standard deviation.

| Cohort Stratum | $n \times K$ | ICC(2,1) | ICC(3,1) | ICC(2,3) | $SD_e$ (pts) | Unanimous < 5.0 ($\kappa$) | Unanimous < 6.0 ($\kappa$) |
|---|---|---|---|---|---|---|---|
| Cohort C: Uncurated Wild Literature | $10 \times 3$ | 0.8583 | 0.8502 | 0.9479 | 0.630 | 80.0% (0.729) | 80.0% (0.659) |
| Cohort A: Retracted / Refuted | $10 \times 3$ | 0.9200 | 0.9330 | 0.9718 | 0.431 | 90.0% (0.712) | 100.0% (1.000) |
| Cohort B: Benchmark Controls | $10 \times 3$ | 0.9126 | 0.9368 | 0.9691 | 0.216 | 100.0% (1.000) | 100.0% (1.000) |
| Pooled Calibration + Wild Sample | $30 \times 3$ | 0.9659 | 0.9678 | 0.9884 | 0.456 | 90.0% (0.864) | 93.3% (0.910) |
| Cross-Cohort Natural Replicates ($r = 0.951$) | $29 \times 2$ | — | — | — | Med $\|\Delta\| = 0.60$ | 89.7% conc. | 89.7% conc. |

Table S3: Blinded evaluation scores and defect rates across all eight contemporary 2020–2025 everyday subfields ($N = 320$, $n = 40$ per field), matched preprint-to-published pairs ($N = 50$ pairs $= 100$ texts), and the balanced open-review concordance cohort ($N = 100$ papers, 261 human referees, 363 substantive human critiques).

| Cohort | Scientific Subfield / Stratum | $n$ | Mean $\pm$ SD | Median (IQR / Core) | < 5.0 (%) | < 6.0 / Recall |
|---|---|---|---|---|---|---|
| In-Domain ($n = 40$/field) | Molecular Evolution | 40 | $6.20 \pm 1.88$ | 6.35 (5.17–7.55) | 20.0% | 45.0% |
| In-Domain ($n = 40$/field) | Viral Evolution | 40 | $6.19 \pm 1.44$ | 6.45 (5.08–7.43) | 25.0% | 37.5% |
| In-Domain ($n = 40$/field) | Cancer & Microbiome (20 Onc. + 20 Host) | 40 | $4.71 \pm 1.67$ | 4.65 (3.45–5.93) | 57.5% | 75.0% |
| In-Domain ($n = 40$/field) | Statistical Genetics | 40 | $4.39 \pm 2.00$ | 4.45 (2.48–5.83) | 60.0% | 75.0% |
| In-Domain ($n = 40$/field) | Transcriptomics & CompBio | 40 | $4.13 \pm 1.76$ | 3.90 (3.02–5.25) | 72.5% | 85.0% |
| In-Domain Total ($N = 200$) | All 5 Core Biomedical Subfields | 200 | $5.12 \pm 1.96$ | 5.20 (3.60–6.60) | 47.0% | 63.5% |
| In-Domain Pilot ($N = 100$) | Initial Cohort C Calibration Slice | 100 | $5.47 \pm 1.78$ | 5.45 (4.10–6.80) | 43.0% | 61.0% |
| OOD ($n = 40$/field) | Neuroscience & Neuroimaging | 40 | $4.80 \pm 1.75$ | 4.65 (3.48–5.65) | 57.5% | 80.0% |
| OOD ($n = 40$/field) | Ecology & Macroevolution | 40 | $4.72 \pm 1.62$ | 4.40 (3.58–5.05) | 67.5% | 80.0% |
| OOD ($n = 40$/field) | Structural Bioinfo & AI Drug | 40 | $3.91 \pm 1.89$ | 3.60 (2.38–5.03) | 72.5% | 87.5% |
| OOD Total ($N = 120$) | All 3 Out-of-Distribution Subfields | 120 | $4.48 \pm 1.79$ | 4.40 (3.30–5.27) | 65.8% | 82.5% |
| All 8 Fields ($N = 320$) | Combined Everyday Cohort (40/field) | 320 | $4.88 \pm 1.92$ | 4.70 (3.40–6.33) | 54.1% | 70.6% |
| Paired Preprint–Pub ($N = 50$) | Selective Flagship/Society ($\Delta Core_8$ / $\Delta Tot_{10}$) | 25 pairs | $5.52 \rightarrow 5.90$ (/8.0) | $\Delta = +0.38$ ($p = 0.019$) / $+0.62$ ($p = 0.0017$) | 20/25 (80%) imp. | 4/25 (16%) declined |
| Paired Preprint–Pub ($N = 50$) | Open-Access Megajournals ($\Delta Core_8$ / $\Delta Tot_{10}$) | 25 pairs | $4.72 \rightarrow 4.90$ (/8.0) | $\Delta = +0.18$ ($p = 0.089$) / $+0.30$ ($p = 0.047$) | 16/25 (64%) imp. | 9/25 (36%) declined |
| Paired Preprint–Pub ($N = 50$) | All Matched Pairs Combined ($\Delta Core_8$ / $\Delta Tot_{10}$) | 50 pairs | $5.12 \rightarrow 5.40$ (/8.0) | $\Delta = +0.28$ ($p = 0.0034$) / $+0.46$ ($p = 0.0002$) | 36/50 (72%) imp. | 12/50 (24%) declined |
| Paired Preprint–Pub ($N = 50$) | Selective vs. Megajournal (ANCOVA / Raw $\Delta\Delta$) | 25 vs. 25 | $+0.50 \pm 0.23$ Tot ($p = 0.033$) | $+0.35 \pm 0.18$ Core ($p = 0.063$) / Raw $+0.20$ ($p = 0.29$) | 80% vs. 64% imp. | 16% vs. 36% declined |
| Open-Review ($N = 100$) | Selective Flagship / Society (135 Referees) | 50 | $7.49 \pm 1.38$ | 7.75 ($Core_8 = 5.87$) | 8.0% | 97.7% (42/43) unres. recall |
| Open-Review ($N = 100$) | Open-Review Megajournal (*PLOS ONE*, 126 Refs) | 50 | $5.01 \pm 1.69$ | 5.10 ($Core_8 = 3.79$) | 48.0% | 98.1% (53/54) unres. recall |
| Open-Review ($N = 100$) | All Open-Review Combined (261 Referees) | 100 | $6.25 \pm 1.98$ | 6.60 ($Core_8 = 4.83$) | 28.0% | 97.9% (95/97) unres. / 78.2% all |
| Open-Review ($N = 100$) | Referee Depth & Anomalies (Sel. vs. Mega) | 50 vs. 50 | 72% vs. 12% Thorough | 0% vs. 38% Superficial | 2% vs. 58% Anomalies | 385 AI-only catches (3.85/paper) |

Table S4: Thirty-five-year secular quality trajectory (1991–2025, $N = 915$ verified full-text articles across seven 5-year windows and four subfield tracks after excluding 85 archival missing-body-text stubs), within-journal fixed-effect and Oaxaca–Blinder decomposition summaries, and structural biomedical covariate partitions across the pooled wild corpus ($N = 1,076$).

| Partition Dimension | Stratum / Window | $n$ | Composite (/10) | $Core_8$ (/8.0) | < 5.0 (%) | < 6.0 (%) |
|---|---|---|---|---|---|---|
| 5-Year Window ($N = 915$) | 1991–1995 | 73 | $7.61 \pm 1.43$ | 6.21 (77.6%) | 6.8% | 13.7% |
| 5-Year Window ($N = 915$) | 1996–2000 | 124 | $7.47 \pm 1.22$ | 6.12 (76.5%) | 4.0% | 10.5% |
| 5-Year Window ($N = 915$) | 2001–2005 | 142 | $6.48 \pm 1.51$ | 5.23 (65.4%) | 18.3% | 31.7% |
| 5-Year Window ($N = 915$) | 2006–2010 | 144 | $6.87 \pm 1.48$ | 5.52 (69.0%) | 11.8% | 25.7% |
| 5-Year Window ($N = 915$) | 2011–2015 | 144 | $6.50 \pm 1.64$ | 5.21 (65.1%) | 20.1% | 34.7% |
| 5-Year Window ($N = 915$) | 2016–2020 | 144 | $5.93 \pm 2.11$ | 4.69 (58.6%) | 36.1% | 47.9% |
| 5-Year Window ($N = 915$) | 2021–2025 | 144 | $5.79 \pm 1.83$ | 4.54 (56.7%) | 31.9% | 52.1% |
| Subfield Track ($N = 915$) | Molecular Evolution & Phylogenetics | 233 | $6.96 \pm 1.55$ | 5.56 ($r = -0.30, p = 3.1 \times 10^{-6}$) | 14.2% | 22.7% |
| Subfield Track ($N = 915$) | Gene Expression & Genomics | 227 | $6.67 \pm 1.57$ | 5.37 ($r = -0.37, p = 1.2 \times 10^{-8}$) | 15.4% | 30.0% |
| Subfield Track ($N = 915$) | Viral Evolution & Virology | 226 | $6.34 \pm 1.98$ | 5.14 ($r = -0.36, p = 2.1 \times 10^{-8}$) | 24.8% | 37.6% |
| Subfield Track ($N = 915$) | Population & Statistical Genetics | 229 | $6.30 \pm 1.80$ | 5.04 ($r = -0.31, p = 2.5 \times 10^{-6}$) | 24.5% | 40.6% |
| Decomposition ($N = 915$) | Within-Journal Fixed Effect (30 Journals) | 549 | $-0.23$ pts/dec | $-0.24$ pts/dec ($p = 0.0074$) | $r = -0.094$ | $p = 0.027$ |
| Decomposition ($N = 915$) | Oaxaca–Blinder (1990s vs. 2020s, $-1.82$ pts) | 427 | 54.1% Compositional | 45.9% Within-Stratum | $-0.98$ pts | $-0.84$ pts |
| Decomposition ($N = 915$) | Equal-Year-Weighted Trend (28 Calendar Yrs) | 915 | $-0.77$ pts/dec | $7.76 \rightarrow 7.12 \rightarrow 6.45 \rightarrow 5.25$ | $r = -0.787$ | $p = 6.8 \times 10^{-7}$ |
| Decomposition ($N = 1,076$) | Multivariable OLS (Decade + Mega + DryLab) | 1,076 | $\beta_{dec} = -0.52$ ($p < 10^{-15}$) | $\beta_{mega} = -0.51$ ($p = 3.0 \times 10^{-6}$) | $\beta_{dry} = -0.71$ | $p = 6.3 \times 10^{-10}$ |
| Venue Model ($N = 1,076$) | Legacy / Society / Selective Journals | 654 | $6.75 \pm 1.79$ | $5.44 \pm 1.53$ | 17.3% | 28.7% |
| Venue Model ($N = 1,076$) | Strict Commercial Open-Access Megajournals | 422 | $5.80 \pm 1.73$ | $4.56 \pm 1.48$ | 33.9% | 51.9% |
| Modality ($N = 1,076$) | Wet-Lab Primary Expt. | 408 | $6.81 \pm 1.68$ | $5.51 \pm 1.45$ | 16.4% | 28.9% |
| Modality ($N = 1,076$) | Hybrid Comp. + Wet-Lab | 351 | $6.56 \pm 1.73$ | $5.21 \pm 1.49$ | 19.9% | 33.0% |
| Modality ($N = 1,076$) | Dry-Lab Theory / Sim. | 87 | $6.33 \pm 1.96$ | $5.05 \pm 1.68$ | 24.1% | 34.5% |
| Modality ($N = 1,076$) | Dry-Lab Secondary DB | 230 | $5.33 \pm 1.78$ | $4.21 \pm 1.51$ | 42.6% | 62.2% |
| Tech. Maturity ($N = 1,076$) | Mid-Stage High-Throughput | 348 | $6.65 \pm 1.65$ | $5.29 \pm 1.42$ | 17.0% | 31.0% |
| Tech. Maturity ($N = 1,076$) | Mature / Classical | 616 | $6.62 \pm 1.71$ | $5.34 \pm 1.47$ | 18.5% | 32.1% |
| Tech. Maturity ($N = 1,076$) | Commodity Template | 112 | $4.16 \pm 1.42$ | $3.16 \pm 1.16$ | 74.1% | 90.2% |
| Epistemic Stage ($N = 1,076$) | Basic Discovery | 679 | $6.89 \pm 1.68$ | $5.53 \pm 1.46$ | 15.3% | 26.7% |
| Epistemic Stage ($N = 1,076$) | Translational / Biomarker | 215 | $5.65 \pm 1.80$ | $4.47 \pm 1.54$ | 36.3% | 54.0% |
| Epistemic Stage ($N = 1,076$) | Clinical / Observational | 182 | $5.33 \pm 1.62$ | $4.23 \pm 1.40$ | 40.7% | 60.4% |

Table S5: Blinded career self-audits across two senior authors ($N = 164$ papers by S.L.K.P., 2004–2026; $N = 74$ by A.N., 2000–2026; 238 reviews across 222 unique papers) by role, alongside the three lowest-scoring lead papers for each author (4.5–6.2/10) and all middle-author papers scoring $< 5.0$ ($n = 8$).

| Role / Stratum | ID (PMCID) | Year | Venue | Score (/10) | Primary Blinded Audit Summary / Defect Identified |
|---|---|---|---|---|---|
| S.L.K.P. First Author ($n = 10$) | Summary | 2005–2023 | 8 Journals | $8.33 \pm 1.25$ | Median 8.65 (IQR 8.43–9.15), 0.0% $<$ 5.0, 10.0% $<$ 6.0 |
| S.L.K.P. Senior Author ($n = 32$) | Summary | 2006–2025 | 18 Journals | $8.19 \pm 1.08$ | Median 8.50 (IQR 7.80–8.93), 0.0% $<$ 5.0, 6.2% $<$ 6.0 |
| S.L.K.P. All Lead/Senior ($n = 42$) | Summary | 2005–2025 | 22 Journals | $8.22 \pm 1.11$ | Median 8.50 (IQR 8.12–9.00), 0.0% $<$ 5.0, 7.1% $<$ 6.0 |
| S.L.K.P. Middle Co-Author ($n = 122$) | Summary | 2004–2026 | 49 Journals | $7.29 \pm 1.39$ | Median 7.70 (IQR 6.50–8.30), 6.6% $<$ 5.0, 16.4% $<$ 6.0 |
| S.L.K.P. All Career Papers ($N = 164$) | Summary | 2004–2026 | 58 Journals | $7.53 \pm 1.38$ | Median 7.85 (IQR 6.70–8.53), 4.9% $<$ 5.0, 14.0% $<$ 6.0 |
| A.N. First Author ($n = 6$) | Summary | 2000–2020 | 4 Journals | $6.97 \pm 1.03$ | Median 6.85 (IQR 6.22–7.55), 0.0% $<$ 5.0, 16.7% $<$ 6.0 |
| A.N. Senior Author ($n = 22$) | Summary | 2005–2025 | 12 Journals | $7.44 \pm 0.94$ | Median 7.50 (IQR 6.93–8.07), 4.5% $<$ 5.0, 4.5% $<$ 6.0 |
| A.N. All Lead/Senior ($n = 28$) | Summary | 2000–2025 | 14 Journals | $7.34 \pm 0.96$ | Median 7.45 (IQR 6.78–8.03), 3.6% $<$ 5.0, 7.1% $<$ 6.0 |
| A.N. Middle Co-Author ($n = 46$) | Summary | 2002–2026 | 25 Journals | $7.88 \pm 0.72$ | Median 7.80 (IQR 7.50–8.47), 0.0% $<$ 5.0, 0.0% $<$ 6.0 |
| A.N. All Career Papers ($N = 74$) | Summary | 2000–2026 | 31 Journals | $7.68 \pm 0.86$ | Median 7.75 (IQR 7.30–8.28), 1.4% $<$ 5.0, 2.7% $<$ 6.0; Lead vs. Middle $\Delta = -0.55$ pts ($p = 0.012$) |
| Joint Co-Authored Papers ($n = 16$) | Independent Blind Runs | 2007–2026 | 9 Journals | 7.82 vs. 7.88 | Cross-batch correlation $r = 0.652$ ($p = 0.0062$), mean difference 0.06 pts; Repro 1.74 (`<body>`) vs. 1.76/2.0 (`<body>`+`<back>`) |
| S.L.K.P. Dry-Lab Only ($n = 116$) | Lead ($n = 39$) vs. Middle ($n = 77$) | 2004–2026 | 2-Way Modality Control | 8.33 vs. 7.49 | $\Delta = +0.84$ pts (Welch $p = 3.4 \times 10^{-4}$); Multivariable $\beta_{\text{lead}} = +0.75$ pts ($p = 0.0025$) |
| S.L.K.P. 4-Way Sub-Modality ($N = 164$) | Theory/Methods ($n = 59$) vs. Sec. DB ($n = 57$) | 2004–2026 | 4-Way Modality Split | 8.40 vs. 7.99 / 7.95 vs. 7.23 | 79% (33/42) of Lead vs. 21% (26/122) of Middle are Methods/Theory; 4-way OLS $\beta_{\text{lead}} = +0.42$ pts ($p = 0.098$) |
| S.L.K.P. Blinding Sensitivity ($N = 148$) | Excl. 16 Residual `github.com/veg` Papers | 2004–2026 | Lead ($n = 32$) vs. Middle ($n = 116$) | 8.26 vs. 7.29 | $\Delta = +0.96$ pts ($p = 5.6 \times 10^{-5}$); Excl. all 109 software-mention papers ($N = 55$): 8.40 vs. 7.08 ($\Delta = +1.32$, $p = 0.0038$) |
| Meta-Audit ($N = 2$ Drafts) | `MANUSCRIPT_META_01` & `02` (This Paper) | 2026 | Blinded Self-Audit | 7.8 | Core 6.2–6.3/8.0 (Rigor 2.6, Claims 1.9, Bio 1.7–1.8, Repro 1.5–1.6): caught untested preprint interaction (ANCOVA $p = 0.033$, pilot raw $p = 0.13$), year-2020 window clustering, 5-vs-6 upstream subset mismatch, and text-vs.-table count/SD discrepancies; all fixed in final text. |
| S.L.K.P. Senior ($<$ 6.0) | `SP_ANON_044` (PMC11289888) | 2024 | *Front Bioinform* | 5.0 | Duplicate Table 2 rows (TN vs. Alberta), impossible node count (1,364 nodes for $N = 295$), unit-scaling errors (100×), and unvalidated heuristic constants (`AUTO-TUNE`). |
| S.L.K.P. First ($<$ 6.0) | `SP_ANON_028` (PMC1480537) | 2006 | *PLOS Comput Biol* | 5.2 | Base-rate fallacy used to justify unadjusted $p \leq 0.05$ across 296 codons (7 sites found vs. $\sim 7.4$ expected under null); core `IFEL` method validated. |
| S.L.K.P. Senior ($<$ 6.0) | `SP_ANON_162` (PMC11228553) | 2024 | *Front Bioinform* | 5.4 | Gap-inflated 627-codon alignment for 340-aa Rem2 misclassifying 251 gap columns as invariant sites; disordered-tail confounding in `AlphaFold2` RMSD; mixed `hg19`/`hg38` table coordinates. |
| A.N. Senior ($<$ 5.0) | `AN_ANON_016` (PMC10288729) | 2023 | *BMC Bioinform* | 4.5 | PDB template leakage in BioGRID cross-validation benchmark and SARS-CoV-2 Spike–MOG receptor prediction driven by antibody-mimicry threading onto neutralizing antibodies (`6XC2`, `7C01`). |
| A.N. First ($<$ 6.0) | `AN_ANON_001` (PMC1186735) | 2005 | *PLOS Genet* | 5.8 | Claims "evolutionary oscillation" between overlapping reading frames (*XLαs/ALEX*) despite no branch ratio differing from 1 ($p = 0.14$–$0.77$) and non-independent shared codon positions. |
| A.N. First (6.2) | `AN_ANON_003` (PMC155263) | 2002 | *Genome Res* | 6.2 | Claims a $\sim 2.6\%$ false-positive rate in the $K_A/K_S$ coding-potential test as an empirical specificity advantage over gene predictors, when 2.6% is a mathematical consequence of testing at $\alpha = 0.05$. |
| Middle Author ($<$ 5.0) | `SP_ANON_103` (PMC9404229) | 2022 | *Environ Res* | 2.2 | Pre-pandemic 2019 Italian SARS-CoV-2 claim: 0/11 RT-qPCR positive, 0/9 $\text{PRNT}_{80}$ positive, single-amplicon nested-PCR carryover of derived 2020 `B.1` control. |
| Middle Author ($<$ 5.0) | `SP_ANON_034` (PMC4592012) | 2015 | *PLOS ONE* | 3.8 | Cross-sectional HIV DNA comparison in CSF vs. blood ($N = 28$) with unadjusted multiple comparisons and over-extrapolated compartmental persistence claims. |
| Middle Author ($<$ 5.0) | `SP_ANON_135` (PMC8927931) | 2022 | *BMJ Glob Health* | 3.8 | Commentary/opinion essay defending early-origin SARS-CoV-2 PCR claims without primary experimental validation or contamination controls. |
| Middle Author ($<$ 5.0) | `SP_ANON_109` (PMC4865637) | 2016 | *Nature* | 4.1 | Ongoing HIV-1 replication under ART claimed from months 0–6 decay window ($N = 3$), 70-cycle nested PCR without UMIs, and negative/conflicting rates in Extended Data Table 2. |
| Middle Author ($<$ 5.0) | `SP_ANON_003` (PMC8135569) | 2021 | *Mol Biol Evol* | 4.2 | Tumor `SCITE` algorithm applied to skewed pandemic surveillance, $>$ 1% filter erasing Wuhan basal diversity, and circular `COI` outgroup flipping (`proCoV2`). |
| Middle Author ($<$ 5.0) | `SP_ANON_021` (PMC8939770) | 2022 | *Sci Transl Med* | 4.5 | Target-predictor autocorrelation in forecasting 3-month SARS-CoV-2 spread from current spread, passenger hitchhiking inflation, and retrospective collection-date slicing. |
| Middle Author ($<$ 5.0) | `SP_ANON_092` (PMC11598840) | 2024 | *Viruses* | 4.6 | Descriptive VSIV outbreak phylogenomics with unvalidated causal adaptation claims and over-interpreted selection scans on small sample size. |
| Middle Author ($<$ 5.0) | `SP_ANON_026` (PMC5177524) | 2017 | *AIDS* | 4.8 | Observational partner-services cohort with severe missing-data attrition and unadjusted confounding between acute stage and network clustering. |

Table S6: Research funding brackets (NIH RePORTER annual grant support across $n = 222$ NIH-funded everyday papers and total acknowledged grants across $N = 1,076$) and journal Open-Access Article Processing Charge tiers across the pooled everyday corpus ($N = 1,076$, and $N = 1,176$ with calibration controls). Acknowledged grants typically fund broad multi-study research programs rather than a single paper. Bottom rows summarize empirical BARS review token and API costs (1,714 unique papers, 1,874 total reviews; $77.48 Batch / $162.10 On-Demand) extrapolated to PubMed Central and PubMed, human peer-review labor benchmarks, and direct APC outlays on failing studies.

| Economic Dimension | Tier / Bracket | $n$ | Cost / Fee Metric | Mean ± SD (/10) | Median (/10) | < 5.0 (%) |
|---|---|---|---|---|---|---|
| NIH Grant Support ($N = 1,076$) | Unfunded / Non-NIH | 854 | $0 NIH | $6.12 \pm 1.83$ | 6.40 | 27.9% |
| NIH Grant Support ($n = 222$ NIH) | Small / Fellowship / R21 (< $250k/yr) | 31 | Med $158,490/yr | $6.93 \pm 1.51$ | 7.40 | 16.1% |
| NIH Grant Support ($n = 222$ NIH) | 1 Standard R01 ($250k–$600k/yr) | 49 | Med $383,323/yr | $7.11 \pm 1.62$ | 7.70 | 10.2% |
| NIH Grant Support ($n = 222$ NIH) | All 1 Grant (< $600k/yr Combined) | 80 | Med $304,562/yr | $7.04 \pm 1.57$ | 7.60 | 12.5% |
| NIH Grant Support ($n = 222$ NIH) | 2–3 NIH Grants ($0.6–$1.5M/yr) | 76 | Med $949,006/yr | $7.63 \pm 1.45$ | 8.00 | 9.2% |
| NIH Grant Support ($n = 222$ NIH) | Mega-Center (> $1.5M/yr) | 66 | Med $3,200,040/yr | $7.46 \pm 1.13$ | 7.80 | 1.5% |
| Total Grants Count ($N = 1,076$) | 0 Acknowledged Grants | 615 | 0 Grants | $6.14 \pm 1.85$ | 6.40 | 27.5% |
| Total Grants Count ($N = 1,076$) | 1 Grant (Single PI) | 216 | 1 Grant | $6.49 \pm 1.72$ | 6.80 | 20.8% |
| Total Grants Count ($N = 1,076$) | 2–3 Grants (Collaborative) | 181 | 2–3 Grants | $6.71 \pm 1.82$ | 7.20 | 20.4% |
| Total Grants Count ($N = 1,076$) | 4–6 Grants (Multi-Lab) | 51 | 4–6 Grants | $7.35 \pm 1.47$ | 7.80 | 7.8% |
| Total Grants Count ($N = 1,076$) | 7+ Grants (Consortium) | 13 | $\geq 7$ Grants | $7.35 \pm 1.36$ | 8.00 | 7.7% |
| Benchmark Modality ($N = 50$) | Cohort B Dry-Lab Math/Software | 14 | Zero Wet-Lab Cost | $8.46 \pm 0.97$ | 8.75 | 0.0% |
| Benchmark Modality ($N = 50$) | Cohort B Wet-Lab Experimental | 36 | Multi-Assay Wet-Lab | $8.71 \pm 0.76$ | 8.90 | 0.0% |
| Journal APC Tier ($N = 1,076$) | Diamond / Free OA ($0–$1.2k) | 28 | Mean APC $0 | $4.99 \pm 2.11$ | 5.25 | 50.0% |
| Journal APC Tier ($N = 1,076$) | Commercial Megajournals & OA ($1.7k–$3.9k, 422 strict + 37 OA) | 459 | Mean APC $2,879 | $5.76 \pm 1.74$ | 5.80 | 35.1% |
| Journal APC Tier ($N = 1,076$) | Academic Society / Univ Press ($2.0k–$4.2k) | 361 | Mean APC $3,242 | $6.48 \pm 1.80$ | 6.60 | 20.5% |
| Journal APC Tier ($N = 1,076$) | High-APC Selective ($4.5k–$7.0k) | 216 | Mean APC $5,579 | $7.62 \pm 1.17$ | 8.00 | 3.2% |
| Journal APC Tier ($N = 1,076$) | High-Fee Flagships ($9.5k–$12.3k) | 12 | Mean APC $11,528 | $7.80 \pm 1.49$ | 8.30 | 0.0% |
| Journal APC Tier ($N = 1,176$ All) | High-Fee Flagships (Everyday + Calibration) | 51 | Mean APC $11,687 | $7.82 \pm 1.75$ | 8.50 | 9.8% |
| AI vs. Human Review | Single-Pass BARS Review (22,119 tokens) | 1,874 | $0.041 Batch / $0.087 On-Dem | 15,392 in / 6,727 out tok | < 3 min | 97.9% unres. recall |
| AI vs. Human Review | 3-Run Consensus BARS Ensemble (ICC = 0.95) | 1,874 | $0.124 Batch / $0.260 On-Dem | 66,357 total tokens | ∼ 3 min | +3.85 AI-only flaws |
| AI vs. Human Review | Human Peer Review (2.61 refs ×5h + 2h ed.) | 100 | $1,505/paper ($100/hr) | 15.05 hrs/paper | 3,213 words | 17.4k–36.4k× AI cost |
| PubMed Extrapolation | 2025 Annual PMC Open-Access Flow | 921,565 | $38,109 Batch / $79,727 OD | 20.38B tokens/yr | $114k 3-Run | 46.9% of PubMed |
| PubMed Extrapolation | 2025 Entire Annual PubMed Flow | 1,965,335 | $81,272 Batch / $170,025 OD | 43.47B tokens/yr | $244k 3-Run | vs. $2.96B human labor |
| PubMed Extrapolation | All-Time PMC Full-Text Stock | 10,900,000 | $450,743 Batch / $942,982 OD | 241.1B tokens | $1.35M 3-Run | 100% PMC archive |
| PubMed Extrapolation | All-Time PubMed Stock (1781–2025) | 39,400,000 | $1.63M Batch / $3.41M OD | 871.5B tokens | $4.89M 3-Run | ≈ 1 NIH R01 grant |
| Downstream Costs | Wild Failing Papers (< 5.0) Direct APC Outlay | 256 | $830k direct APCs in sample | Mean APC $3,242 | — | Direct publication fees |

Table S7: Blinded evaluation of `gemini-4-argon` (`Gemini 3.1 Pro`) against the 12 models evaluated by Brian et al. (2026) across the first three expanded 2026 *Nature Medicine* clinical trials (`T01`–`T03`; 3 error-seeded and 3 control reviews per model, an illustrative $n = 3$ cross-check) and across all 58 trials in Brian et al. (2026). Net Youden's $J = \text{Sens}_{\text{Maj}} + \text{Spec}_{\text{Ctrl}} - 1$. Models marked with † in Brian et al. (`Opus 5.5`, `Fable 5`, `Opus 5`) were evaluated on only $n = 3$ initial trials (`N`, `I`, `M`) and omitted from `T01-T60`.

| Model | `T01-T03` Major Sens. (%) | `T01-T03` Control FPR (%) | `T01-T03` Specificity (%) | `T01-T03` Youden's $J$ | 58-Trial Total Sens. / Spec. (%) | Calibrated Rigor Score (Clean vs. Seeded) |
|---|---|---|---|---|---|---|
| `gemini-4-argon` **(Ours)** | **84.0%** (7.56/9) | **2.4%** (0.33/14) | **97.6%** | **+0.816** | — | **$6.93 \pm 0.61$ (PASS) vs. $3.00 \pm 0.50$ (FAILED)** |
| `GPT-6.1 Sol` | 92.6% (8.33/9) | 13.7% (1.92/14) | 86.3% | +0.789 | 90.2% / 74.8% | — |
| `Sonnet 5` | 84.3% (7.58/9) | 10.1% (1.42/14) | 89.9% | +0.742 | 81.2% / 82.9% | — |
| `GPT-5.4` | 81.5% (7.33/9) | 7.7% (1.08/14) | 92.3% | +0.738 | 70.9% / 88.2% | — |
| `GPT-6 Luna` | 85.2% (7.67/9) | 14.9% (2.08/14) | 85.1% | +0.703 | 86.3% / 79.4% | — |
| `GPT-5.5` | 81.5% (7.33/9) | 13.1% (1.83/14) | 86.9% | +0.684 | 73.1% / 86.2% | — |
| `Kimi K3` | 81.5% (7.33/9) | 16.1% (2.25/14) | 83.9% | +0.654 | 76.2% / 77.7% | — |
| `GPT-5.4 mini` | 70.4% (6.33/9) | 5.4% (0.75/14) | 94.6% | +0.650 | 59.0% / 90.8% | — |
| `DeepSeek V4 Pro` | 75.9% (6.83/9) | 11.9% (1.67/14) | 88.1% | +0.640 | 68.1% / 86.4% | — |
| `Gemini 3.8 Flash` | 77.8% (7.00/9) | 16.1% (2.25/14) | 83.9% | +0.617 | 66.2% / 84.0% | — |
| `Qwen 3.8 27B` | 74.1% (6.67/9) | 15.5% (2.17/14) | 84.5% | +0.586 | 62.6% / 81.8% | — |
| `gpt-oss-120b` | 59.3% (5.33/9) | 3.6% (0.50/14) | 96.4% | +0.557 | 56.8% / 92.8% | — |
| `Haiku 4.5` | 55.6% (5.00/9) | 10.1% (1.42/14) | 89.9% | +0.455 | 45.8% / 89.1% | — |
| `Opus 5.5`† ($n = 3$ trials only) | — | — | — | — | 92.9% / 71.4% | — |
| `Fable 5`† ($n = 3$ trials only) | — | — | — | — | 92.5% / 74.6% | — |
| `Opus 5`† ($n = 3$ trials only) | — | — | — | — | 89.7% / 79.4% | — |
